# Fast and accurate shared segment detection and relatedness estimation in un-phased genetic data using TRUFFLE

**DOI:** 10.1101/460915

**Authors:** Apostolos Dimitromanolakis, Andrew D. Paterson, Lei Sun

## Abstract

Relationship estimation and segment detection between individuals is an important aspect of disease gene mapping. Existing methods are either tailored for computational efficiency, or require phasing to improve accuracy. We developed TRUFFLE, a method that integrates computational techniques and statistical principles for the identification and visualization of identity-by-descent (IBD) segments using un-phased data. By skipping the haplotype phasing step and, instead, relying on a simpler region-based approach, our method is computationally efficient while maintaining inferential accuracy. In addition, an error model corrects for segment break-ups that occur as a consequence of genotyping errors. TRUFFLE can estimate relatedness for 3.1 million pairs from the 1000 Genomes Project data in a few minutes on a typical laptop computer. Consistent with expectation, we identified only three second cousin or closer pairs across different populations, while commonly used methods identified a large number of such pairs. Similarly, within populations, we identified much fewer related pairs. Compared to methods relying on phased data, TRUFFLE has comparable accuracy but is drastically faster and has fewer broken segments. We also identified specific local genomic regions that are commonly shared within populations, suggesting selection. When applied to pedigree data, we observed 99.6% accuracy in detecting 1^st^ to 5^th^ degree relationships. As genomic datasets become much larger, TRUFFLE can enable disease gene mapping through implicit shared haplotypes by accurate IBD segment detection.

## Introduction

Estimating relatedness and co-ancestry among pairs of individuals is a commonly encountered task in genetic studies. Traditionally, likelihood-based methods (e.g. PREST^1; 2^) or method-of-moments estimators (e.g. PLINK^3^) were used in linkage or association studies, respectively. KING^4^ introduced a computationally efficient kinship estimation approach (KING-kinship) that does not explicitly require allele frequency estimation and presumably could be more robustly applied to relationship inference in non-homogeneous population samples. The method is widely used for the inference of *close* relationships in large-scale genetic data, although it can have a higher error rate for distantly related pairs^5^; also see Results below.

The availability of increasing marker densities in studies using genotyping arrays or sequencing technologies makes a different class of methods that perform identical-by-descent (IBD) segment detection more attractive. These methods estimate recent shared ancestry between pairs of individuals by identifying shared chromosomal segments, and they have been implemented in software such as GERMLINE^6^, BEAGLE Refined IBD^7^, and fastIBD^8; 9^. However, although these methods can provide more refined estimates of shared ancestry, identify long-distance relationships and assist disease mapping, they typically require orders of magnitude more computational time and need accurate phasing of the input data. The resulting application complexities prevent their broader use in large-scale genetic studies.

Methods for IBD segment detection in un-phased data have been proposed, including IBDSeq^10^, Parente2^11^, and the recent shared segment method implemented in the KING software (KING-segment)^12^. Among those, IBDSeq and Parente2 are not fast enough for application to large studies and do not provide genome-wide IBD estimation in a single step. The accuracy profile of the KING-segment method has yet to be reported.

We developed TRUFFLE, a practical method that enables faster and yet accurate identification of IBD1 and IBD2 segments shared between individuals, calculation of averaged IBD sharing probabilities across the genome (or kinship coefficients), and visualization of shared segments using un-phased genetic data. By skipping the haplotype phasing step and, instead, relying on a simpler region-based approach, the proposed method is less computationally intensive and much easier to apply. In addition, a built-in error model corrects for segment break-ups that can occur as a consequence of genotyping errors. Finally, an integrated variant filtering allows direct application of TRUFFLE to raw variant calls from VCF files, without the need for external linkage disequilibrium (LD) pruning of markers.

## Materials and Methods

Without loss of generality, let us consider a single chromosome on which we identify IBD1 and IBD2 segments for a pair of individuals (*a*, *b*) based on available, un-phased bi-allelic single nucleotide polymorphisms (SNPs) data. As in convention, the markers are arranged by physical position, and the genotypes of individual *a* at marker *j* are *G^a^*(*j*) = {0, 1, 2}, where *a* ranges from 1 to *n*, and *j* ranges from 1 to *m*. For every pair of individuals (*a*, *b*), we define

*IBS_2_^a,b^*(*j*): = 1 if *G^a^*(*j*) *= G^b^*(*j*), and = 0 otherwise,
*IBS_12_^a,b^*(*j*): = 1 if abs(*G^a^*(*j*) - *G^b^*(*j*)) < 2, and = 0 otherwise.

*IBS_2_^a,b^*(*j*) tracks if the genotypes at marker *j* are identical between the two individuals, i.e. two alleles shared identity-by-state (IBS), IBS2, while *IBS_12_^a,b^*(*j*) tracks IBS *at least* one. If either of the genotypes at SNP *j* is missing, then both values are defined to be 1 to prevent segment break-up and keep the same set of markers for all pairs analyzed. It is assumed that long continuous stretches of missing data do not exist in datasets that underwent standard quality control.

For each marker *j* on the chromosome we also define *p.IBS2*(*j*) and *p.IBS12*(*j*) to be the expected probability of having, respectively, two alleles and at least one allele shared IBS between a pair of unrelated individuals. These probabilities depend on the minor allele frequency (MAF) of the bi-allelic marker, *maf_j_*, such that *p.IBS2*(*j*) = (*maf_j_*)^4^ + (1-*maf_j_*)^4^ + (2*maf_j_* (1-*maf_j_*))^2^ and *p.IBS12*(*j*) = 1 - 2(*maf_j_*)^2^(1-*maf_j_*)^2^. We can then define *p_2_* and *p_12_*, respectively, as the averaged *p.IBS2*(*j*) and *p.IBS12*(*j*) across all markers. The values of *p_2_* and *p_12_* would in turn depend on the distribution of the MAFs for the panel of bi-allelic markers used. However, we note that the method is generally robust to mis-specification of MAFs (see Results).

### Base model with no genotype error consideration

As a basic model we consider scanning a pair of individuals for long stretches of IBS2 or IBS12. A stretch of multiple consecutive IBS markers are likely to be IBD if (1) it is unlikely to occur by chance, and (2) it extends beyond the LD of the region that is being considered.

For criterion (1), in a typical whole-genome dataset (either by genotyping or sequencing, e.g. the 1000 Genomes Project^13^) and considering only common variants (MAF > 5%, defined globally), the average *p.IBS2*(*j*) over approximately 1,000 markers from a randomly selected region ranges from 0.46 to 0.51, with a mean of *p_2_* = 0.48 over the whole genome (Supplementary Data – Section 2). A consecutive stretch of *k* independent markers that are all IBS2 has a probability of occurring by chance of approximately 0.48*^k^*. Thus, when ignoring LD between markers we could set a length threshold *l2* for declaring IBD2 segments, such that *p_2_^l2^* < 10^−8^. The detection of IBD1 segments through long stretches of IBS sharing is harder, as the probability of at least one allele shared IBS (i.e. IBS12) by chance is substantially higher than IBS2 alone. For example, the average *p.IBS12*(*j*) in a randomly selected region ranges from 0.92 to 0.94, with a mean of *p_12_=0.93* (Supplementary Data – Section 2). To establish a low probability of a false positive for IBD1, as before, we set the minimum length *l1* (typically substantially greater than *l2*) such that *p_2_^l1^*< 10^−8^ for independent markers.

For criterion (2), ideally, using a model that takes into account local LD structure can guide the selection of the minimum segment length required for a particular region. However, LD-based hidden Markov models (HMMs) pose a serious computational burden and are typically thousands of times slower than non-LD models^5^. The need for an accurate and high-resolution genetic map also limits their applicability to individuals of mostly European descent. To reduce the effects of LD without incurring a significant computational time penalty, we consider a basic pruning approach such that markers closer than a specific number of base pairs are removed. We performed sensitivity analysis of the minimum length parameters, *l1* and *l2*, to identify the cutoff values for robust estimation of overall IBD1 and IBD2 sharing (Supplementary Data – Section 3). As a default, segments shorter than 5 Mb for IBD1 or 2 Mb for IBD2 are not considered, although these cutoffs can be adapted using command line options. Filtering of segments by genetic distance in centiMorgan (cM) can also be done, with a set of post-processing scripts. For our analyses here, we have used the genetic map on build GRCh37 as provided in the BEAGLE^8^ website (see Web Resources). Although these default parameter values can be revised by the user, we have found that in practice (see Results below) they work well for datasets with a variety of ancestral compositions, variant densities, and sequencing or array-based platforms.

### Model with genotyping error

A common problem in segment detection is the presence of genotyping error, which breaks apart segments and can easily cause false negatives in segment detection. In addition, *de novo* mutation events can generate spurious errors that will further increase the rate of segment break-ups. For example, with an error rate of 0.5%, two individuals on average will have at least 25 markers with genotyping error in a segment of 5,000 markers. Previous analysis showed that methods accounting for genotyping error like IBDseq have better performance than methods that do not, like Refined IBD^7^ (Figure 2 of ^10^). The error model implemented in TRUFFLE was developed to cope with error rates up to 1%, which might be the case in low-depth sequencing data. In its essence, the proposed approach is a finite deterministic state space model with an unbounded number of states (Figure 1).

**Figure 1:**
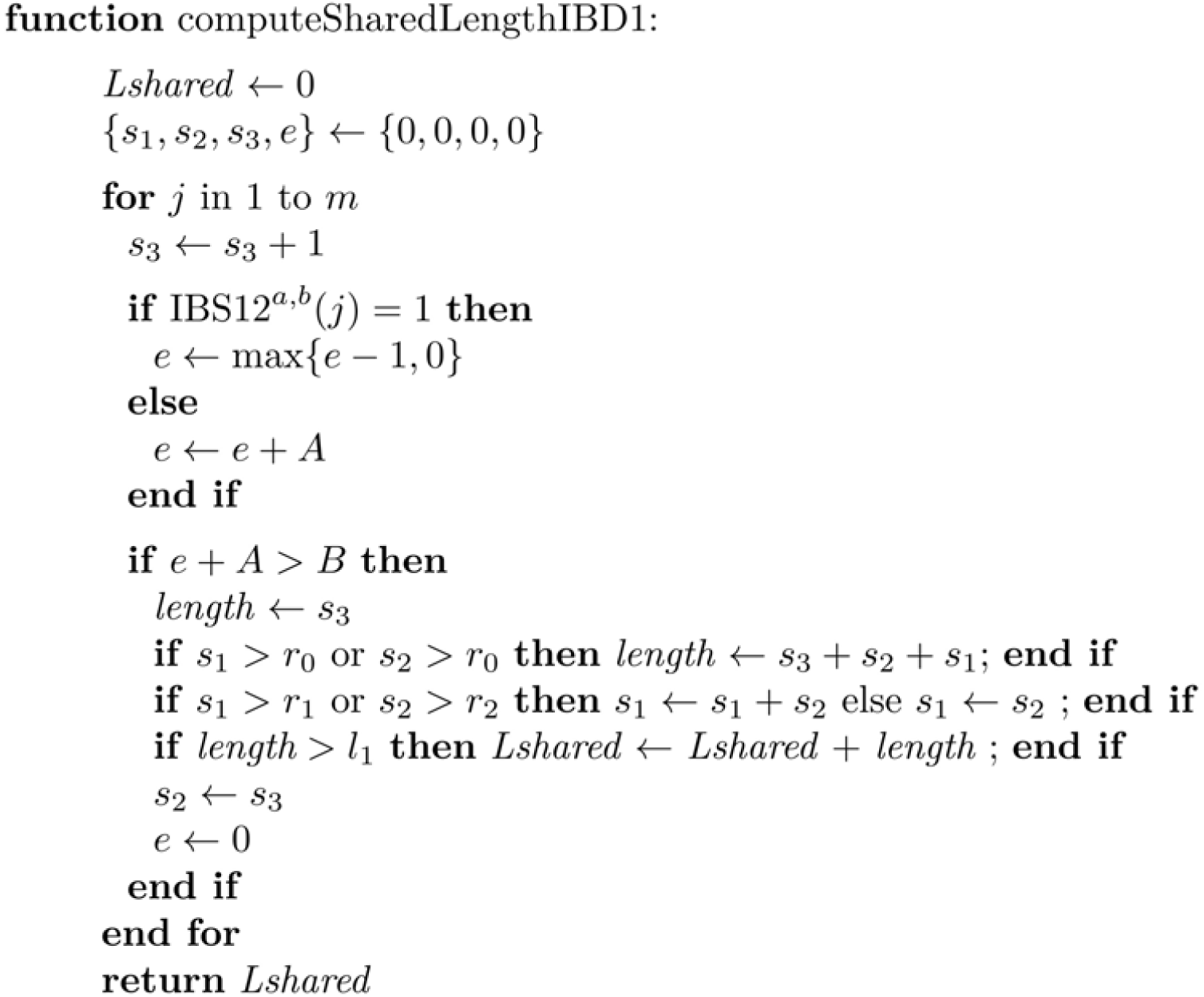
The TRUFFLE algorithm for IBD1 detection with error model. For IBD2 replace *IBS_12_^a,b^*(*j*) with *IBS_2_^a,b^*(*j*).

**Figure 2:**
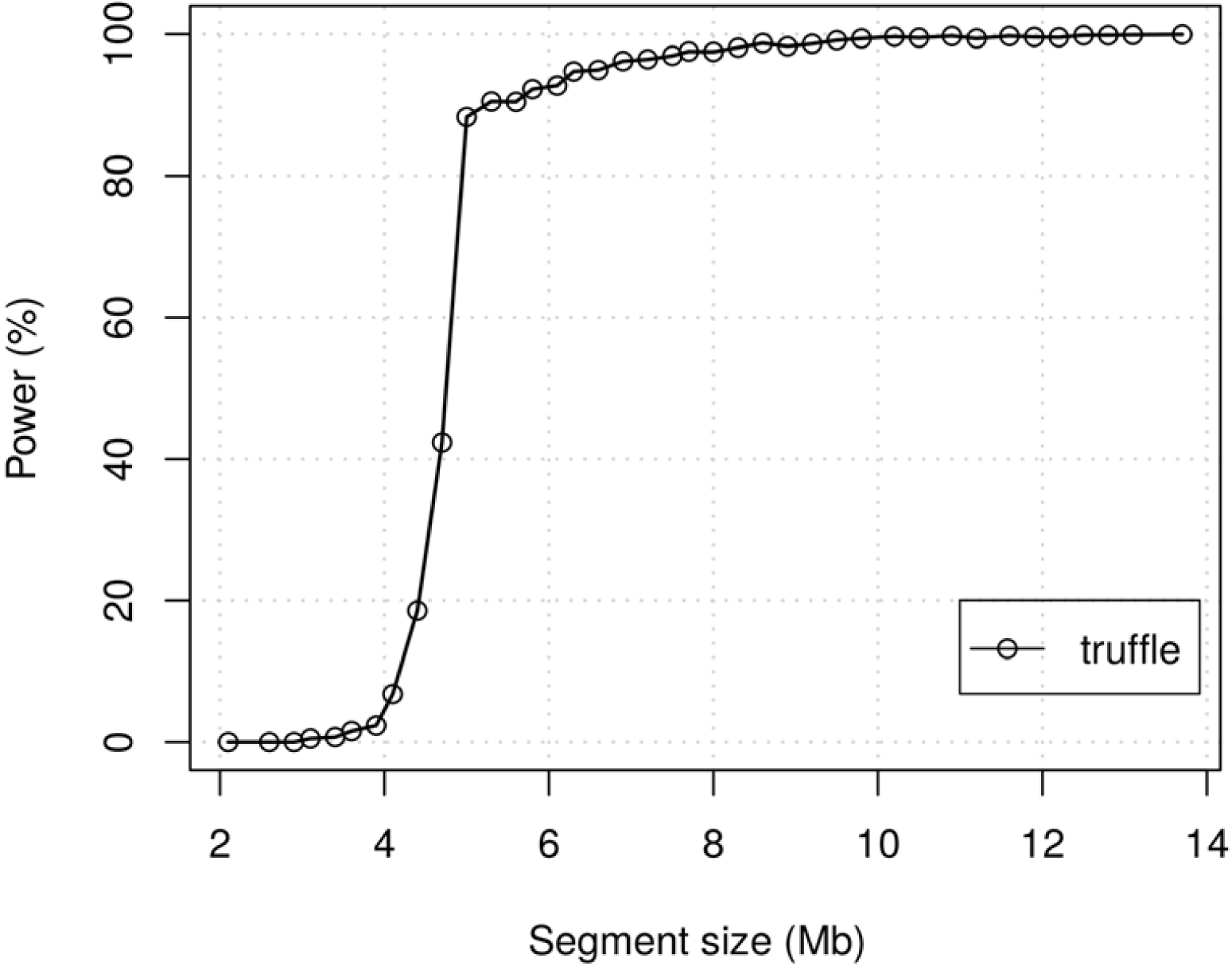
Power of IBD 1 segment detection (α = 4. 6 · 10^−4^) by TRUFFLE stratified by segment size. True shared segments (IBD1) of varying sizes were inserted in simulated variant data (38,174 markers) using simuPOP and the simuGWAS scripts.

For the case of identifying IBD1 segments for a pair of individuals (*a*, *b*), the genome is scanned sequentially from marker 1 to *m*. A set of four states is kept at each marker position: *S_j_* = {*s_1_, s_2_, s_3_, e*}, *j* = 1, …, *m*, with the initial values being *S_0_*= {0, 0, 0, 0}. Intuitively, these four states correspond to the lengths of the last three IBS1 segments (*s_1_, s_2_, s_3_*) that were found and an error load (*e*) of the currently considered segment. A description of the algorithm for identifying IBD1 segments is shown in Figure 1. The algorithm for IBD2 segment detection is identical, but with different values for the five tuning parameters: *A*, *B*, *r_0_*, *r_1_*, and *r_2_*.

The parameter setting of A = 1, B = 0 and r_0_, r_1_, r_2_ = ∞ corresponds to a no genotyping error model. In contrast, given a non-zero *B*, approximately *B/A* errors are allowed for a shared segment before it is considered broken, while short segments are joined together if at least one of them is long enough. In practice, this model is computationally efficient as no complex mathematical operations are required.

The default values for parameters A, B, r_0_, r_1_, and r_2_ are different for detecting IBD1 and IBD2, and they were optimized by simulations. Briefly, for IBD1 we simulated regions containing 20,000 independent variants, with an average probability of IBS12 of 0.93 to 0.95 (representative of typical genotyping datasets, for example see Supplementary Figure S1). Artificial IBD shared segments of 500 markers were added by copying one or two haplotypes of the region from one individual to another; this segment size corresponds to a proportion of IBD1 of 0.125% in a 400k-marker panel. An exhaustive search for the best parameter values was then performed, selecting the ones that maximized the detection power at a false positive error rate of 0.001. For the case of IBD2, similar datasets were simulated, with the markers having an average probability of IBS2 of 0.5, and artificial IBD2 segments of sizes 1 to 5 Mb were added.

### Segment visualization

A significant benefit of IBD segment detections algorithms is that they can provide the locations of IBD1 and IBD2 segments, across the genome. TRUFFLE aids the visualization of such segments by including a set of scripts to create interactive images showing the chromosomes and shared segments; see Results below.

### Implementation

TRUFFLE was implemented in C++. It is readily applicable to genome-wide datasets with high marker densities, even on typical laptop computers. Support for parallel execution in multi-core computers and variant filtering (e.g. MAF, missing rate, and minimal distance between markers in base pairs) is integrated. The input file is a multi-sample VCF file generated ideally from joint calling across samples^14^ and contains all or some autosomes. The input file can be phased, although it is not treated differently. If necessary, users can define parameter values for the minimum segment detection length and the reporting threshold for related pairs. TRUFFLE is available free for non-commercial use (see Web Resources).

## Results

### Power study

To better understand the statistical properties of TRUFFLE to identify shared segments across distant relatives, we pursued simulations using simuPOP v1.1.3 ^15^, following the simulation design of ^16^; the exact simulation scripts are provided in the Supplementary Data.

A single chromosome was simulated, with 38,174 bi-allelic markers having MAF > 5%. The simulation used the HapMap phase III populations TSI and LWK as the initial population composition^3^, simulating a heterogeneous dataset of 3,000 individuals with equal numbers from the two populations. Artificial IBD1 segments of varying sizes were then injected into pairs of individuals within each population.

Under the null condition we set the false positive rate to be 4.6×10^−4^; note that this error rate depends on the parameter values used in the simulation. Although this rate appears to be small, it allows for 2,070 false positives for the 3,000 individuals analyzed because there were about 4.5×10^6^ pairs in total.

For each dataset simulated under the alternative, 100 individuals were randomly selected and 100 artificial IBD1 segments, of lengths ranging from 2 to 14 Mb, were created by copying these 100 segments into another 100 randomly selected individuals. In addition, genotype errors based on an error rate of 0.9% were added to the shared segments. In total, 15,000 datasets were simulated. While TRUFFLE accurately detects large segments (power > 80% for segments > 5 Mb), it has lower power (< 5%) for segments < 4 Mb (Figure 2).

### 1000 Genomes Project data

We applied TRUFFLE to the 20130502 release of the 1000 Genomes phase III data ^13^. The dataset consists of variant calls for 2,504 individuals from 26 populations (five super-populations: 661 Africans, 347 Admixed Americans, 504 East Asians, 489 South Asians, and 503 Europeans). The total number of variants is approximately 88 M before any filtering is applied. These variants were derived from a combination of low and high coverage whole genome sequencing data, high coverage exome sequence, and genome-wide association study (GWAS) array data from two platforms^13^.

For our analyses and method evaluations, three subsets of the bi-allelic markers were generated. The first dataset (A) mirrors what is typically used for relatedness estimation by selecting bi-allelic markers with global MAF > 10% across all populations, and performing LD pruning using PLINK v1.90b3.44 ^3^; the indep-pairwise procedure with parameter values 2000, 200, and 0.1 for the number of markers in window, shift, and r^2^ criteria. A total of 63,126 markers remained in dataset (A). The second dataset (B) was derived by selecting markers with MAF > 5% and with a spacing of at least 5 kb between two consecutive markers, resulting in 469,470 markers remaining. Dataset (B) was generated to evaluate the performance of TRUFFLE when the computationally expensive step of LD-pruning is avoided. Unlike dataset (A), dataset (B) can be internally generated by TRUFFLE in a single step from a multi-sample VCF file, streamlining the cryptic relatedness analysis for whole-genome sequencing studies. In addition, due to the higher marker density dataset (B) allows for detection of shorter shared segments. Finally, dataset (C) included all ∼12M biallelic SNPs with MAF>1%. Variants with missingness >2% were excluded for all 3 datasets.

For comparison, datasets (A) and (B) were analyzed using the two different approaches implemented in KING version 2.1.6, i.e. KING-kinship ^4^, and the more recent KING-segment.

Despite over 3 M pairs of individuals to be examined, the running time of TRUFFLE was 1.6 mins, using 8 cores of a 2Ghz Xeon CPU processor for dataset (A). The running time of the KING-segment procedure was only 9.7 sec but came at the cost of robustness; see below. TRUFFLE running time for dataset (B) was 9.7 mins because of the increased number of markers, however it does not require the LD-pruning step that involved 115 mins of computing time using the indep-pairwise procedure in PLINK. KING time for dataset (B) was 40 secs.

For relationship estimation, KING identified a significant number of distant relationship pairs and appeared to be quite sensitive to the density of markers, in contrast with TRUFFLE (Table 1, Supplementary Figures S5 and S6). With a kinship coefficient cutoff of > 0.0097 for declaring second cousin or closer relatedness, the KING-kinship method reported 573,326 pairs while the KING-segment method reported 214 pairs using the low-density LD-pruned marker dataset (A). When using the higher-density bp-pruned dataset (B), KING-segment also reported an unusually high number of 28,012 second cousin or closer related pairs, among which 14,079 pairs are across populations (Table 1). In contrast, TRUFFLE estimated 200 pairs with kinship coefficient equal or greater to second cousin using (A) and 229 using (B), among which 189 pairs are overlapping and only two and three pairs are across populations.

**Table 1:** Comparison of relationship estimation in 2,504 individuals from the 1000 Genomes. **Top:** Kinship estimation using TRUFFLE v1.38 and KING v2.1.6 for the two datasets (A) and (B) generated from the 1000 Genomes Project phase 3 data^13^. The numbers of pairs that fall under four different kinship cutoffs are shown. **Bottom:** The corresponding numbers of pairs where the two individuals belong to two different populations are also shown. Large numbers of such pairs are more likely to be false positives. ^1^ Cutoff chosen as the midpoint between the kinship of the relationship in consideration and the kinship of the next more distant relationship considered in this table. Pairs counted have the estimated kinship greater than the specified cutoff for each row. KING results for dataset (C) (∼12M SNPs with MAF>1%) are provided for comparison since this is recommended for KING.

|  |  | Dataset (A) |  |  | Dataset (B) |  |  | Dataset (C) |
| --- | --- | --- | --- | --- | --- | --- | --- | --- |
| Relationship | Cutoff <sup>1</sup> | TRUFFLE | KING kinship | KING segment | TRUFFLE | KING kinship | KING segment | KING segment |
| All pairs |  |  |  |  |  |  |  |  |
| Full sibling or parent-offspring | 0.1875 | 12 | 12 | 12 | 12 | 12 | 12 | 12 |
| First cousin or closer | 0.035 | 61 | 14201 | 59 | 61 | 90 | 1726 | 81 |
| Second cousin or closer | 0.0097 | 200 | 573326 | 214 | 229 | 34927 | 28012 | 2036 |
| Third cousin or closer | 0.0024 | 1543 | 684657 | 1976 | 2815 | 172800 | 35799 | 8180 |
| Pairs observed across populations |  |  |  |  |  |  |  |  |
| Full sibling or parent-offspring | 0.1875 | 1 | 1 | 1 | 1 | 1 | 1 | 1 |
| First cousin or closer | 0.035 | 2 | 6524 | 2 | 2 | 8 | 211 | 2 |
| Second cousin or closer | 0.0097 | 2 | 467149 | 2 | 3 | 16746 | 14079 | 21 |
| Third cousin or closer | 0.0024 | 25 | 573081 | 30 | 244 | 110274 | 18704 | 251 |

For first-degree relative pairs, results of all three methods (KING-kinship or segment, and TRUFFLE) agree: there were four full-sib pairs and eight parent-offspring pairs reported (Table 1). Looking closer, the estimated IBD2 sharing by the KING-segment method are 18.3%, 12.3%, 14.8%, and 17.3%, respectively, for the identified four full-sib pairs using dataset (A), with a mean of 15.7%. This is noticeably different from the mean value of 25.1% using dataset (B) also based on KING-segment. In contrast, the mean values based on TRUFFLE are, 25.6% and 25.9%, using respectively datasets (A) and (B).

We also analyzed dataset (C) using KING, as recommended by KING. The results are indeed improved compared to using dataset (B) for KING (see Table 1). However, using this many markers negates any computational advantage of KING over TRUFFLE as now the running time for KING was 29 minutes, about 3 times slower than TRUFFLE. Nevertheless, both KING and TRUFFLE are much faster than phased methods. For example, an earlier numerical experiment showed that BEAGLE Refined IBD and GERMLINE required 64 CPU days for phasing a dataset with ∼2,500 individuals and ∼500,000 SNPs, compared to 5 minutes when using an earlier, slower KING method ^5^

Our primary analyses used MAFs estimated from all available individuals, which are simple to implement when analyzing cross-population pairs. To study the effect of using globally defined MAFs on the TRUFFLE analyses for individual populations, we re-analyzed dataset (B) (∼470,000 SNPs with global MAF >5%). We re-screened the SNPs requiring the MAF > 5% in the CEU sample alone (∼430,000 SNPs), then performed the TRUFFLE analysis again in CEU. IBD segment estimates are very similar between the two analyses (correlation > 0.99). This is true for another population, the LWK African sample, analyzed in a similar way. In addition, the location of shared segments generally were not sensitive to the number of markers did not vary when either global or population-specific MAFs were used (Supplementary Figure S12).

Visualization of the exact locations of detected IBD segments can be generated using TRUFFLE post-processing scripts. Figure 3 illustrates segment locations for two selected related pairs from the 1000 Genomes Project, obtained in dataset (B).

**Figure 3:**
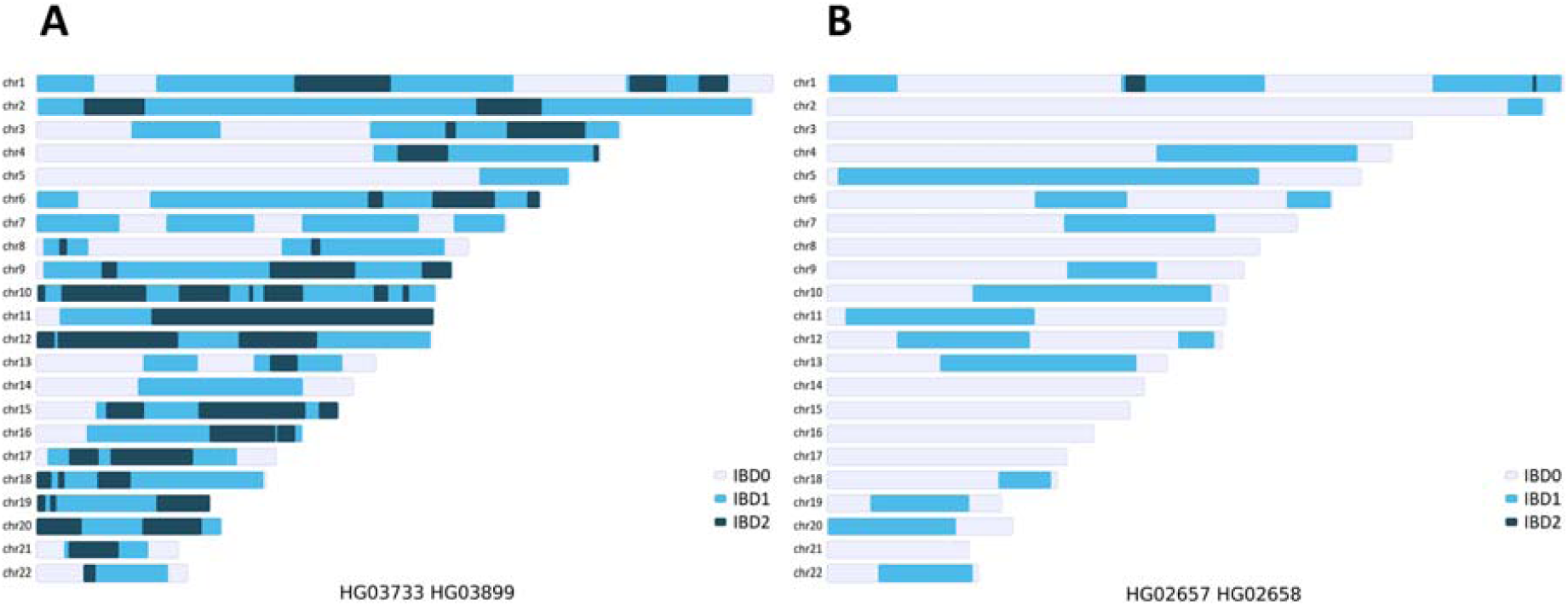
Locations of shared segments identified by TRUFFLE in two pairs from the 1000 Genomes data. A. A putatively full-sib pair from the STU population showing numerous *IBD1* and *IBD2* shared segments. B. A putatively more distant related pair from the PJL population with estimated *IBD1* of 32% and *IBD2* of 0.48% across the genome.

### Short shared segment analysis

Genomic sharing among unrelated individuals is common and has been described previously. For example, analyses of HapMap II data revealed patterns of segment sharing among seemingly unrelated individuals^17^. It was estimated that, on average, any two individuals from the same population share approximately 0.5% of their genome through recent IBD, and 10% to 30% of the pairs share at least one region of their genome IBD.

We performed a scan of all the 2,504 individuals from the 1000 Genomes Project for IBD1 and IBD2 segments using TRUFFLE and dataset (B). To this end, a minimum length of 1,000 markers was used as a cutoff to detect both IBD1 and IBD2 segments; this is different from the earlier default recommendations of 5 Mb for IBD1 and 2 Mb for IBD. To understand the characteristics of locally shared IBD segments between apparently unrelated individuals, we removed segments shared between 574 pairs that are closely related (estimated average IBD1+IBD2 > 0.02). Among the remaining pairs, the minimum segment length detected was 5.45 Mb for IBD1 and 5.54 Mb for IBD2, and the maximum lengths were, respectively, 68.0 Mb (pair HG00641-HG01162 within the PUR population) for IBD1 and 11.9 Mb (pair HG02348-HG01967 within the PEL population, Peruvians from Lima, Peru). In total, there were 956,577 IBD1 segments and 575 IBD2 segments.

Greater than 30% of the pairs in the Puerto Ricans from Puerto Rico population (PUR) share at least 0.5% of their genomes IBD 1 (Figure 4, panel (a)) and 62% of pairs share at least one segment of length > 10 cM (Supplementary Figure S9). The sharing is even more extensive for segments > 5 cM, where more than 82% of pairs share at least one segment. These findings align with previous analyses using Refined IBD of BEAGLE^7^; for example ^13^ showed that Puerto Ricans have one of the lowest effective population sizes. The average sharing length in the PUR population was 28.5 cM among all pairs; for comparison, the average length in CEU, GBR, and TSI was 2.37, 4.28, and 3.62 cM respectively.

**Figure 4:**
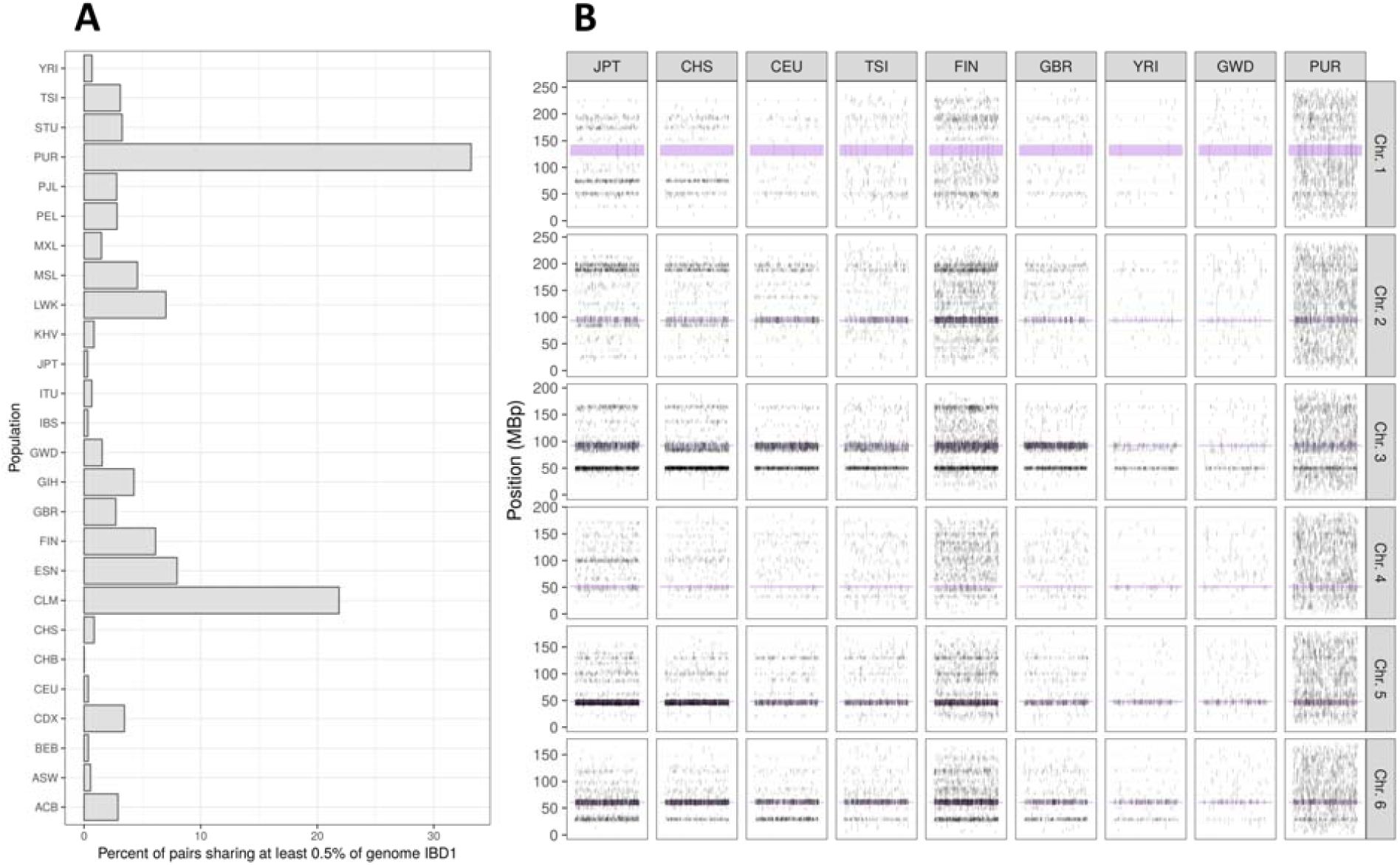
Shared segments among the 1000 Genomes populations. A. Proportion of pairs within a population that share at least 0.5% of their genome as IBD1. B. Distribution of segment locations identified within pairs of the same population for the first six chromosomes and nine selected populations. The segments are positioned randomly on the x-axis to aid visualization and reduce over-plotting. The centromere location is denoted with a purple segment

The Finnish in Finland sample (FIN) also showed extensive segment sharing (average length 16.1 cM), to an extent higher than the other three European populations (CEU, TSI, and GBR). More than 18% of the pairs in the FIN population share a segment of length >10 cM, in contrast to 0.7% for CEU.

Distribution of segments in panel (b) of Figure 4 (and supplementary Figure S10) also suggests sharing hotspots across the genome, likely due to reduced recombination rates in those locations. Similar to our analyses, other approaches have found and reported such hotspots^18; 19^. A high proportion of the identified segments fall in specific genomic regions across multiple different populations (e.g. CEU and CHS in Figure 4); see Figure S8 for all 26 populations. Some of the hotspot regions match centromeres of specific chromosomes, indicative of reduced recombination rates at those regions but perhaps also low SNP density. These patterns are less pronounced in African populations (e.g. GWD and YRI), possibly reflecting their higher genetic diversity^20^. Such IBD hotspots shared across populations could inflate relationship estimation and exclusion of hotspots could alter the interpretation of distant relationship estimation.

### Comparison with genotyping array data

To assess the applicability of TRUFFLE to genotyping array data, we used individuals genotyped on the Illumina Omni2.5 array as part of the 1000 Genomes Project ^13^. Quality control has been previously performed^21^, and the post-quality control data were downloaded from the TCAG website, consisting of 2,318 individuals and 1,989,184 SNPs.

To mirror the dataset (B) generated from the 1000 Genomes combined sequencing and array data, we applied TRUFFLE to 322,849 bi-allelic markers with MAF > 5% and having minimum distance of 5 kb between markers. Among the 2,318 individuals in the array dataset, 1,693 were common with those in dataset (B). Thus, we compared the kinship estimates for all the pairs involving those 1,693 individuals.

The correlation of TRUFFLE kinship estimates, using array or combined sequencing and array data, was very high with a sample correlation of 0.998 for pairs estimated as having kinship coefficient > 0.01 in either of the two datasets. Essentially, the inference of relatives closer than third cousin is identical between array and sequencing data. Among all pairs, the sample correlation was 0.932 (Supplementary data – Figure S8), with a mean difference in kinship estimates of 2.9×10^−4^ (standard error of 8.2×10^−4^).

### Comparison of total lengths of shared segments from TRUFFLE and KING with previously published BEAGLE Refined IBD results in the 1000 Genomes

The Refined IBD procedure in BEAGLE ^7^ is a hidden HMM approach for detecting IBD segments that accounts for LD structure in phased genotype data. Previously, shared segment analysis using BEAGLE version 4.1 was conducted for the 1000 Genomes phase III data and reported by the 1000 Genomes Project ^13^. In their analysis, bi-allelic SNPs with more than 10 copies of the minor allele were used and results were post-processed to delete small gaps between segments (as Refined IBD does not directly account for genotyping error). We compared the reported results with those of TRUFFLE, as well as with the estimates from the KING-segment method applied to the dataset (A) or (B) as previously. The BEAGLE Refined IBD shared segment results were obtained from the 1000 Genomes project ftp site (see Web Resources). Because the reported segments in BEAGLE did not distinguish between IBD1 and IBD2, we compared the total length of all IBD segments in each individual pair, which is proportional to the estimated *p.IBD1*+*p.IBD2*.

The agreement between the Refined BEAGLE IBD segment estimation and TRUFFLE is very high with a sample correlation of 0.956 for dataset (A) (Figure S7) and 0.966 for dataset (B) (Figure 5). In contrast, the correlation of BEAGLE with KING-segment was 0.971 for dataset (A) (Figure S7) and 0.355 for (B), consistent with the over-estimation of distant relatedness for dataset (B) as seen in Table 1. Using dataset (C), the correlation between KING and BEAGLE results was 0.88 (Figure 5). Essentially, TRUFFLE is a compromise between statistical and computational efficiency. KING is faster, but using the results inferred from BEAGLE Refined IBD procedure as benchmarks, TRUFFLE provides a better approximation of the shared segment lengths than KING. BEAGLE is more accurate but TRUFFLE does not need haplotype phasing that is computationally costly, or a detailed genetic map that may not be available for a population of interest.

**Figure 5:**
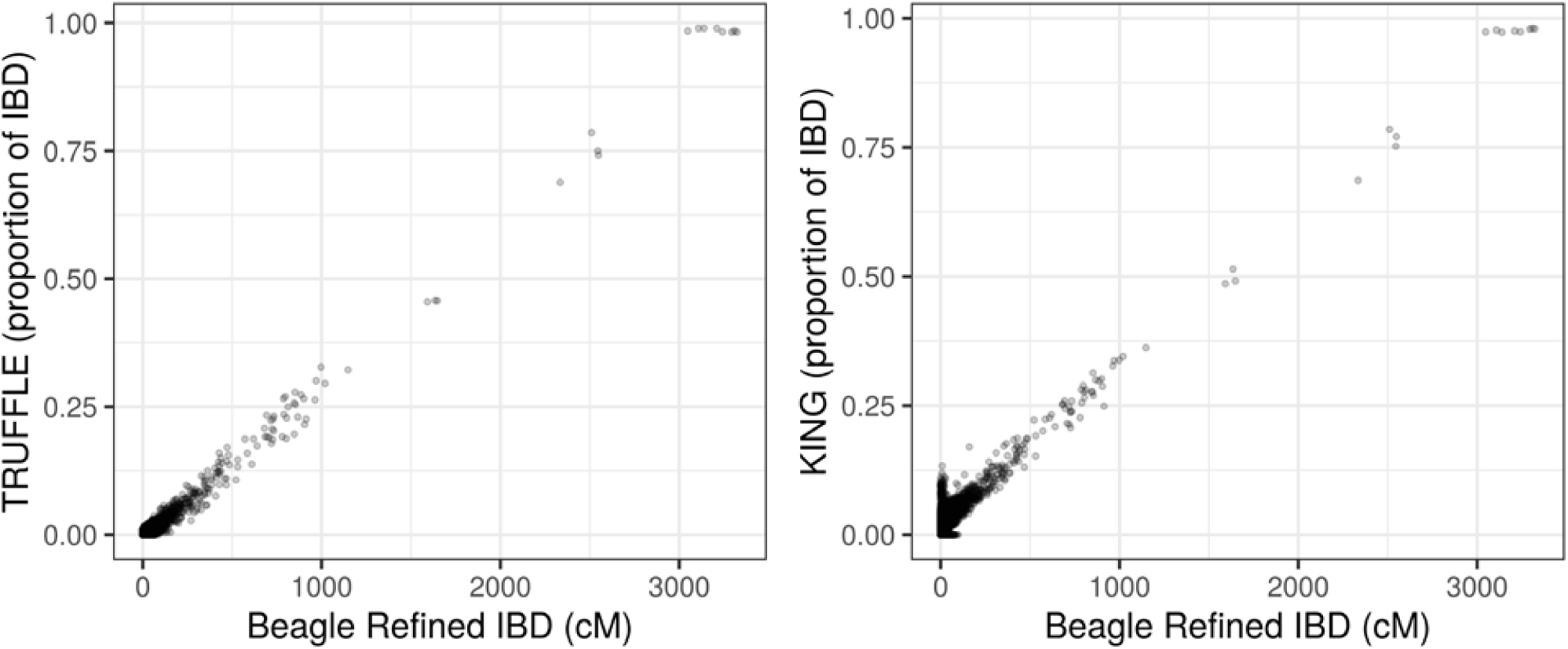
Comparison of total shared segment lengths identified in the 1000 Genomes by TRUFFLE and KING to BEAGLE Refined IBD. The left figure for TRUFFLE vs. BEAGLE using dataset (B), and the right figure for KING-segment vs. BEAGLE using dataset (C) as recommended by KING. BEAGLE results were downloaded from the 1000 Genomes project ftp site. Because BEAGLE did not distinguish between IBD1 and IBD2, the Y-axis shows the estimated *p.IBD1*+*p.IBD2* by TRUFFLE or KING for comparison. We did not convert the cM segment sizes to Mb for BEAGLE in the x-axis, as it would require population specific genetic maps.

### Comparison of locations of shared segments from TRUFFLE with BEAGLE Refined IBD, GERMLINE and KING in the 1000 Genomes

To compare the specific locations of shared segments between pairs of individuals we focused on chromosome 1 data from both dataset (B) (32,926 SNPs) and dataset (C) (943,790 SNPs) and compared 4 methods, including two (Refined IBD ^7^ and GERMLINE ^6^) that require phased input. Here we used the previously phased data from the 1000 Genomes analysis group using both BEAGLE and SHAPEIT2. Specifically we ran: GERMLINE (Web Resources) using the options: -bits 32 -haploid -min_m 3 -err_hom 4 -err_het 1. (We also ran GERMLINE using the default option of -bits 128, but there were excessive segment breakups for the parent-offspring pairs.); BEAGLE Refined IBD segment detection method using the default options, with the genetic map provided with BEAGLE (Web Resources); KING ^4^ v 2.1.6 using the --ibdseg method for inferring segment locations; TRUFFLE v1.38 using the default options. For Refined IBD we present the results both before and after merging segments using the merge-ibd-segments.26Feb19.29e.jar program with the recommended options (Web Resources).

In the absence of de novo mutations, either from single variants or large indels or CNVs, we expect parent-offspring pairs to have IBD1 across the autosomes: this represents a reasonable gold-standard. Therefore Figure 6 shows results for all eight parent-offspring pairs. We have also selected a random pair from other more distant relationships for comparisons of lengths and positions of identified IBD segments (Supplementary Figure S11).

**Figure 6:**
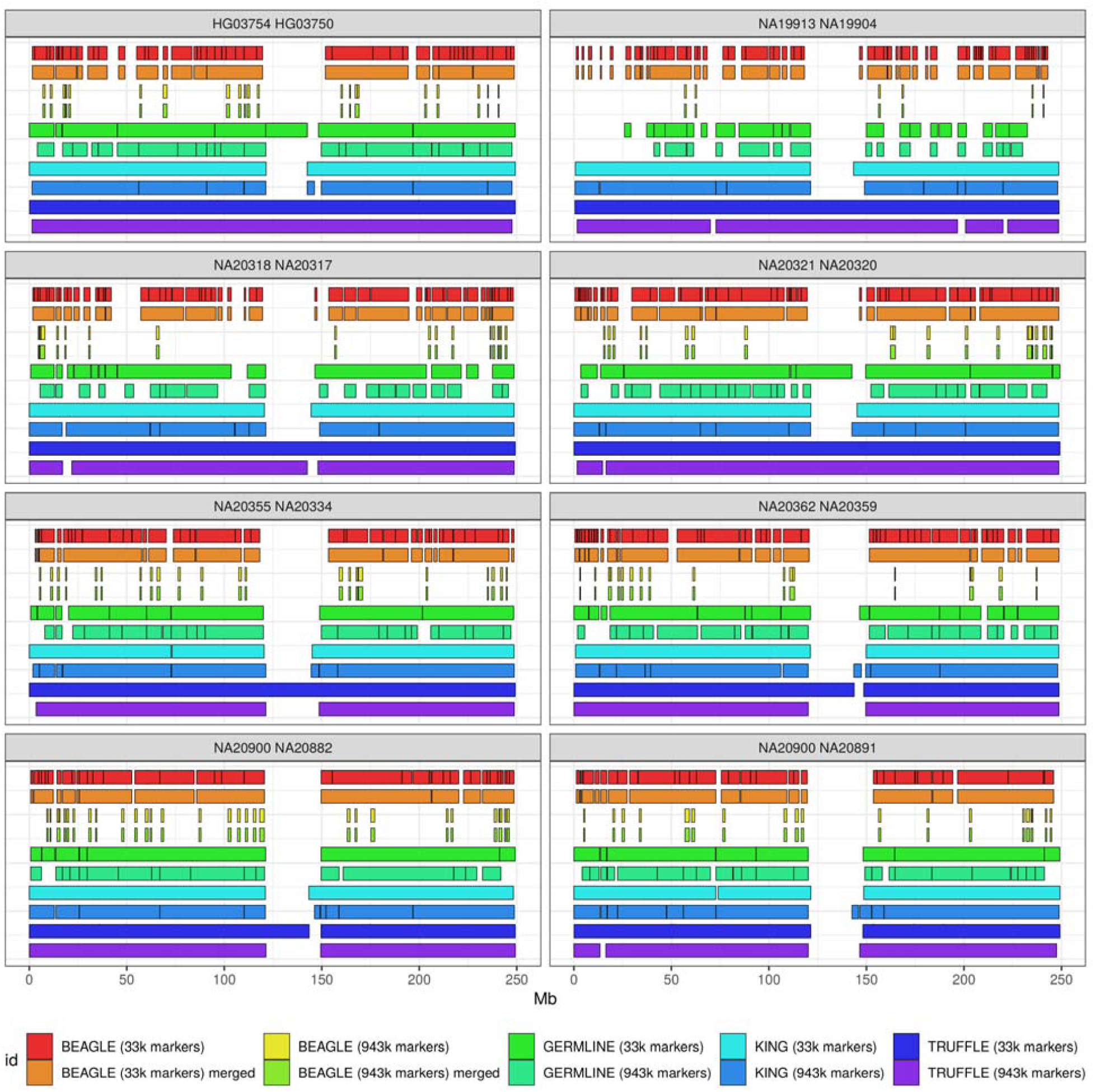
Comparison of locations of IBD segments on chromosome 1 from the 1000 Genomes Project for 8 parent-offspring pairs using different methods and variant densities. The data are from phase 3 release 5. KING and TRUFFLE can work on un-phased data, and BEAGLE Refined IBD and GERMLINE were applied to the data previously phased by the 1000 Genomes analysis group using both BEAGLE and Shapeit2. In the absence of *de novo* mutations we expect parent-offspring pairs to have IBD1 across the autosomes: representing a gold-standard. The 33k SNPs have MAF > 5% with > 5 kb between two consecutive SNPs with missing rate < 2%, and the 943k SNPs have MAF > 1% and missing rate < 2%. Positions are based on build 37, where the centromere is located at 121.5 - 142.5 Mb.

We conclude that TRUFFLE generally identifies segments of expected lengths (i.e. whole chromosome for parent-offspring pairs), does not have segments broken up, and is relatively robust to the selection of markers in comparison to most of the other methods. In contrast, the two methods that require phased data, BEAGLE Refined IBD and GERMLINE, show many short segments. Note, most methods do not identify IBD at the centromere of chromosome 1 due to low marker density across this large region (> 20 Mb).

Consistent with expectations, using 943k chromosome 1 SNPs with MAF > 1% typically produces more segment breaks, likely due to the fact that the genotyping error rate of some of these variants is higher than the 33k SNPs which all have MAF > 5% ^22^. Similar results are observed for other types of relative pairs, including randomly selected full-sibs, first cousins, along with more distantly related pairs (Figure S11).

### Estimation of accuracy in pedigree data

We analyzed Affymetrix 6.0 array data from 822 genotyped individuals from 173 pedigrees. The data were part of the Genetic Analysis Workshop 20 (GAW20) project, and provided by the Genetics of Lipid Lowering Drugs and Diet Network (GOLDN) study ^23; 24^. The GOLDN study recruited European American pedigrees with at least two siblings from the communities of Minneapolis, MN, and Salt Lake City. The average pedigree size was 17.8 individuals, with an average of 4.75 genotyped individuals per pedigree. The numbers of reported relationship pairs within the pedigrees are shown in supplementary Table S1. Individuals from different pedigrees are presumed to be unrelated.

As part of the GAW20 data release, 718,542 autosomal bi-allelic SNPs were available for analysis. We applied TRUFFLE to a reduced variant set of 210,181 markers, selected as having MAF > 5% and minimum distance of 5 kb between two consecutive markers. The TRUFFLE analysis of 337,431 pairs required 32 seconds on a Core-i7 desktop computer (including both across and within pedigree pairs).

Overall, the TRUFFLE kinship estimates closely matched the reported relationships (Figure 7), even using this non-LD pruned variant subset. Overall, 99.6% of the relationships were estimated correctly to within one degree, where the estimated degree of relationship is computed from the estimated kinship, 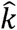, as the closest integer to − log_2_ 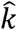 − 1 (Supplementary Table S1).

**Figure 7:**
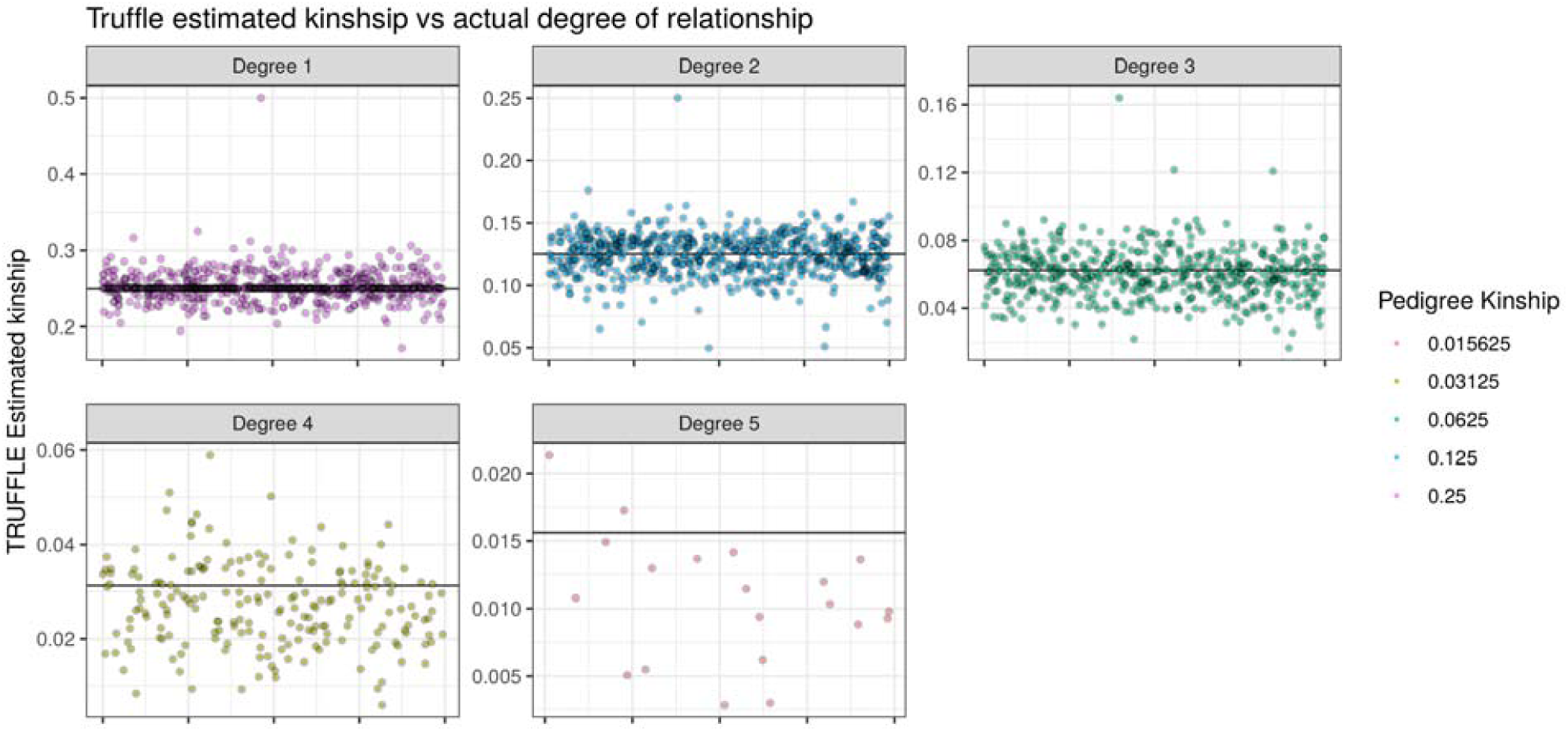
Kinship estimation in the GAW20 data. Only relative pairs, as specified in the study pedigree, are shown. Results are grouped by the degree of relationship based on the given pedigree structure, and the horizontal line shows the expected kinship coefficient for each panel. With each group, pairs are randomly ordered on the x-axis.

Even though the estimated relationship was in line with the specified one overall, 11 pairs of 4^th^ degree or closer related individuals appeared to have mis-specified relationship, with a ratio of reported vs. estimated kinship greater than 2 (or ½). In addition, 13 pairs showed strong evidence of inbreeding, having estimated *p.IBD2* > 2%. Among the 684 presumed unrelated within-pedigree pairs, the average estimated kinship was 0.0013. However, among the 334,431 between-pedigree presumably unrelated pairs, 30 showed estimated relationship of 5^th^ degree or closer. These individuals share 4.3% to 11.7% of their genome as IBD1, with shared segments occurring in multiple locations across their genomes and an average of 6.2 shared segments per pair; they are likely to be true relatives.

For comparison, we also applied KING to the GOLDN data. For first and second degree relatives, the TRUFFLE and KING kinship estimates are consistent with each other, and with the pedigree-based values. For more distantly related relatives, while TRUFFLE slightly underestimates the relationship, KING slightly overestimates (Supplementary Table S2).

## Discussion

In applications to population-based data^13^ and family-based pedigree data^23^, TRUFFLE provides accurate IBD1 and IBD2 estimation within a few minutes of computer time for a complete scan of all pairs in a sample using un-phased genome-wide data. Although it is likely that HMM based models, such as Refined IBD ^7^, will ultimately have more power in detecting short (1-3 Mb) segments, their computational burden and requirement for phased data prohibits their widespread use.

Our power and pedigree studies showed that TRUFFLE has high accuracy in providing pedigree relationship estimation and distinguishing distant cousin pairs sharing > 5 Mb segments (corresponding to a putative 10^th^ degree relative pair). Our applications also demonstrated TRUFFLE’s applicability to both sequencing and array-based studies. The visualization of the exact locations of detected IBD segments is another useful feature of TRUFFLE. Compared to other commonly used methods, TRUFFLE appears to suffer less from breaking up segments (Figure 6).

Although it is easy to apply TRUFFLE to studies with up to 20,000 individuals, further enhancements and speed improvements would be needed to make application to large-scale, population-based genetic studies routine. When analyzing > 20k individuals with > 500k variants there could be memory issues with the current TRUFFLE implementation. Based on the empirical evidence from analyzing dataset (A) (∼50k variants) and dataset (B) (∼500k variants) in the 1000 Genomes Project, we also recommend reducing the number of variants used as an initial screening step, or analyzing each chromosome separately as practical mitigating solutions. Hashing and dictionary-based approaches are useful future directions by means of avoiding the all-pairs quadratic number of comparisons. Although such methods have been previously applied to segment detection in phased data^6^, application of such methods to un-phased data is not trivial and would require new algorithmic techniques and inferential methods.

Common variants are more informative for IBD inference than rare variants. Genotype accuracy declines with lower MAF, particularly for variants derived from low-coverage NGS ^22^. Future work will focus on rare variants, including having error models that differ by MAF and depth.

The relatedness from the X-chromosome can be wildly different from the autosomes, as it follows a different inheritance pattern. Because of the lower recombination rate^25^, the X-chromosome will require different models for the analysis and discovery of shared segments. The pseudo-autosomal regions of the X-chromosome (PAR1-3) would also require specific handling, which is of future research interest.

Overall, TRUFFLE provides a significant improvement in the applicability of IBD segment detection methods to many types of genetic studies. The combination of ease of use, accurate IBD estimation for both distant and close relationships, and segment location visualization greatly extend the goal of traditional relationship inference methods. TRUFFLE can enable disease mapping and population genetics through implicit shared haplotypes by accurate IBD segment detection focusing on overlapping segments from multiple pairs of affected individuals.

## Supplemental Data

Supplemental data include 12 figures, 2 tables and accompanying text.

## Acknowledgements

The authors have no conflict of interest to declare.

We sincerely thank the two reviewers, the Associate Editor and the Editor for constructive comments that substantially improved the manuscript. We also thank the investigators of the GOLDN project and the Genetic Analysis Workshop advisory committee for permission to use GAW20 data. This research was funded by the Canadian Institutes of Health Research (CIHR, MOP-310732-G-CEAA-117978) and the Natural Sciences and Engineering Research Council of Canada (NSERC, RGPIN-250053) to LS.

## Web Resources

TRUFFLE v1.38: https://adimitromanolakis.github.io/truffle-website/

PLINK v1.90b3.44: https://www.cog-genomics.org/plink2

KING v2.1.6: http://people.virginia.edu/~wc9c/KING/

1000 Genomes VCF: http://ftp.1000genomes.ebi.ac.uk/vol1/ftp/release/20130502/

1000 Genome array data: http://www.tcag.ca/tools/1000genomes.html

Human Genetic Maps GRCh37: http://bochet.gcc.biostat.washington.edu/beagle/genetic_maps/

BEAGLE Refined IBD Results: http://ftp.1000genomes.ebi.ac.uk/vol1/ftp/release/20130502/supporting/ibd_by_pair/

GERMLINE version 1.5.3 released on 06/10/2018; http://gusevlab.org/software/germline/

BEAGLE Refined IBD version released on February 26, 2019; http://faculty.washington.edu/browning/refined-ibd.html

