## Supplementary data for "Fast and accurate shared segment detection and relatedness estimation in un-phased genetic data using TRUFFLE"

### 1. Description of downloaded data from 1000 genomes

The 1000 genomes dataset 20130502 was downloaded as 22 VCF files for the 22 autosomal chromosomes, from the following url:

ftp://ftp.1000genomes.ebi.ac.uk/vol1/ftp/release/20130502/.

Filtering and LD-pruning of variants for the generation of dataset A was performed using PLINK (version 1.90b44).

The resulting vcf files are available for download at the github URL:

https://github.com/adimitromanolakis/truffle

### 2. Average allele sharing in 1000 genomes dataset

Compute average probability of sharing 1,2 or 2 alleles is important in establishing a baseline minimum region size for called segments. For dataset B with 469k autosomal markers, we computes the probability of sharing 1 or 2 alleles (1-p.IBS0) in windows of 1000 markers across the genome (figure S1), which ranged from 0.916 to 0.937 ( mean 0.926 ). For p.IBS2, the mean value across the genome for 1000 marker windows ranged from 0.459 to 0.511 (mean 0.483).

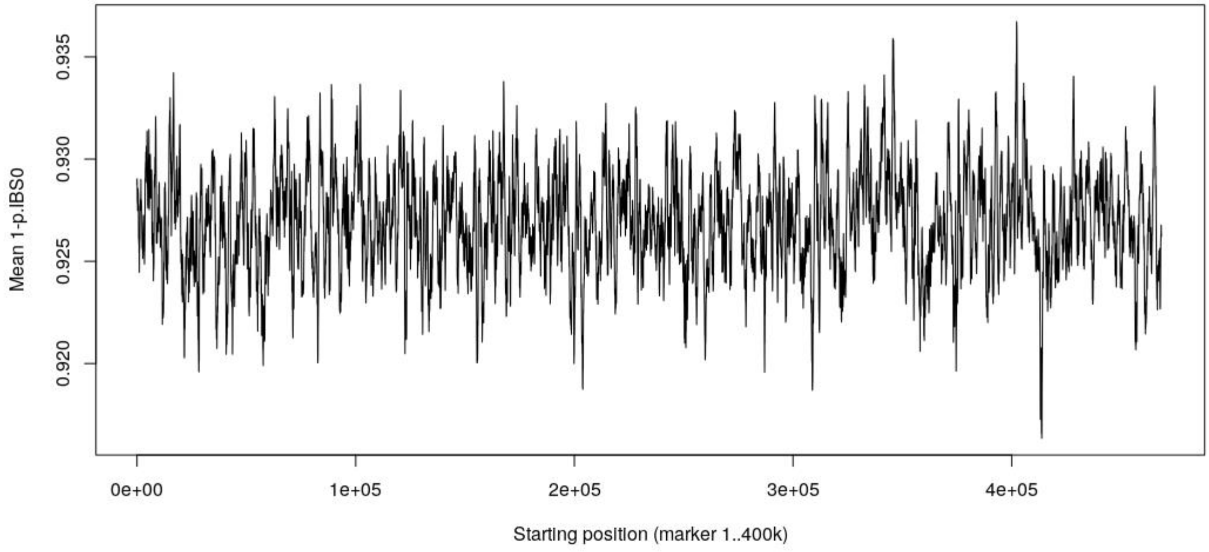


**Figure S1:** Probability of sharing 1 or 2 alleles (averaged over all pairs of individuals) at every genomic location. 1000 Genomes data – Dataset B

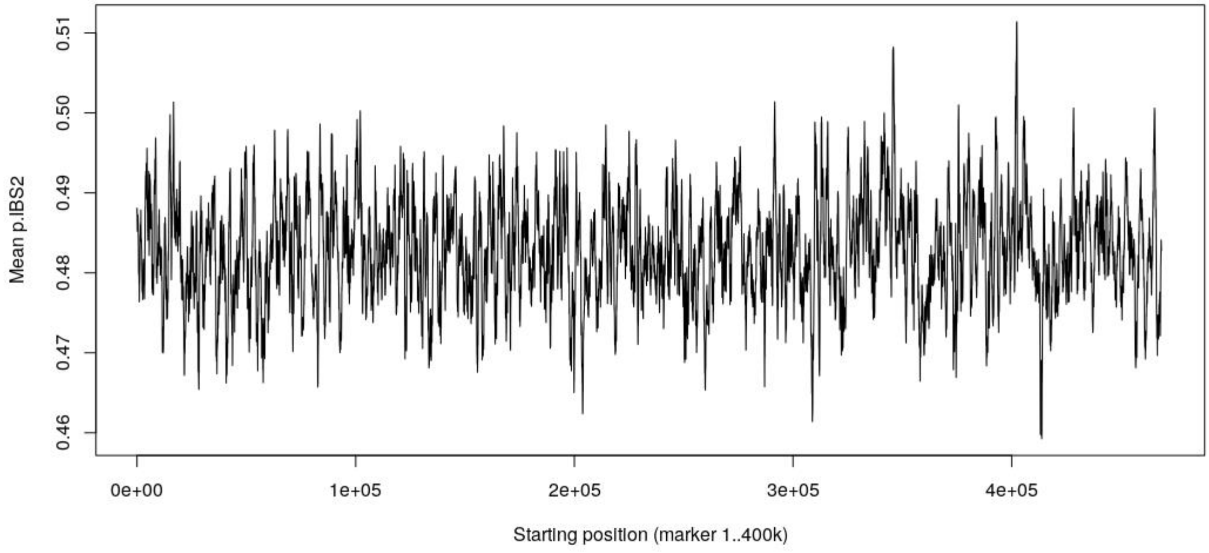


**Figure S2:** Probability of sharing 2 alleles (averaged over all pairs of individuals) at every genomic location. 1000 Genomes data – Dataset B

### 3. Sensitivity analysis of the segment length cutoff parameter

We analyzed the sensitivity of the IBD1 and IBD2 estimates, by varying the estimation window parameter S from 0.1 to 4, and analyzing all pairwise relationships for 47 individuals from dataset B. These 47 individuals included individuals previously identified by TRUFFLE as parent-offspring, full-sibling. In addition we included individuals being identified as unrelated (as pairs with very low identified IBD). The variant set analyzed was dataset B.

Internally, truffle computes a minimum accepted segment length of an IBS1 or 2 segment for its inclusion as IBD. The parameter ***S*** in TRUFFLE specifies the adjustment factor of this length. For example a value of 2, will specify that only segments twice as long as the default value will be accepted. The trajectories of IBD1 and IBD2 estimates of those pairs (figure S3), show how the corresponding relatedness estimation varies by adjusting ***S***. Increasing values of S reduce the estimates of IBD1 and IBD2 as smaller segments are not counted. Decreasing values of S have the opposite effect, by included very small IBS1 and IBS2, which occur likely by chance or because of the LD between the markers.

For a small number of pairs, the trajectories of 2 specific pairs (selected as 1 parent-offspring, 1 full sibling) are compared to the model without the provision for genotyping error (figure S4).

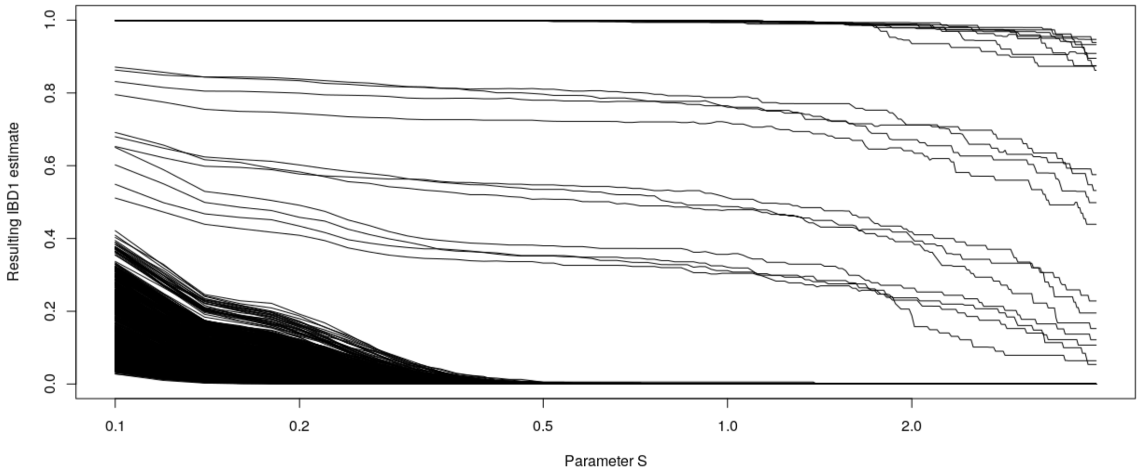


(a)

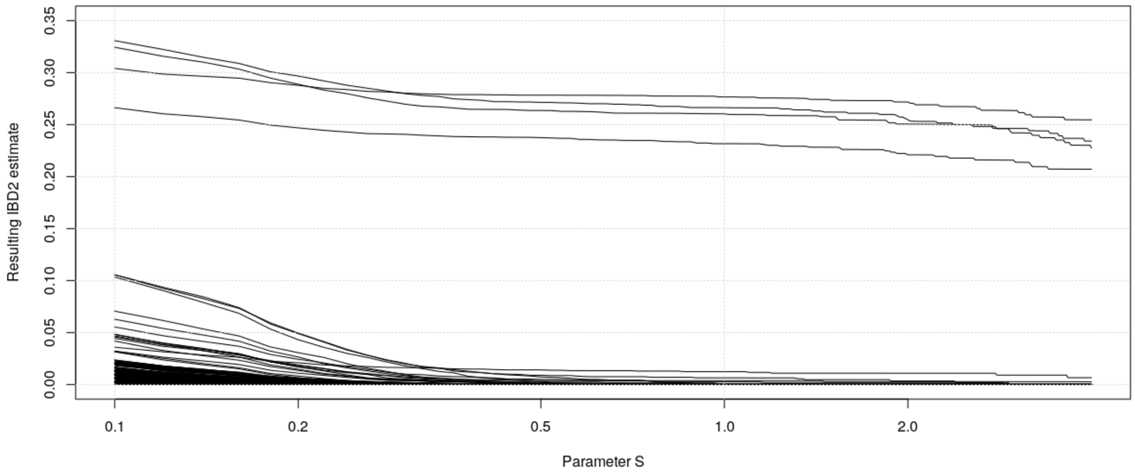


(b)

**Figure S3:** Trajectories of IBD1 and IBD2 estimates for 1081 pairs from the 1000 genomes data, by varying the parameter L in truffle. The pairs included the 12 identified 1st degree relative pairs, 7 additional pairs of 2nd to 3rd degree relatedness and a number of randomly selected pairs with low detected IBD. Each line represents one pair of individuals and the estimation of IBD1 or 2 for all values of the parameter S. (a) IBD1 estimation vs L. (b) IBD2 estimation vs L.

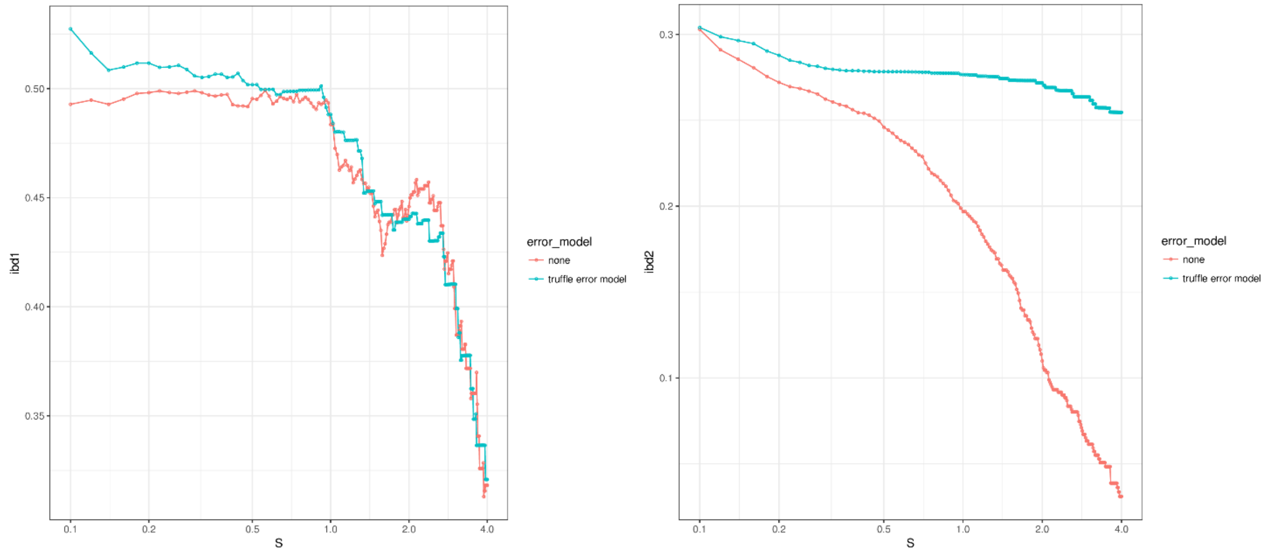


*(a) Full Sibling Pair:*

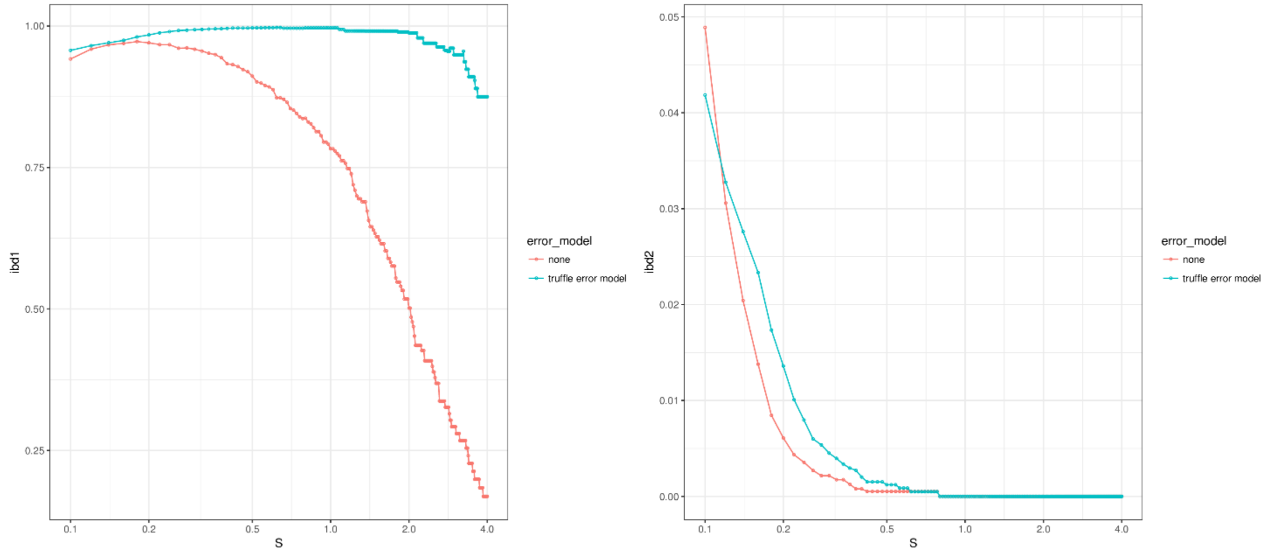


(b) Parent offspring pair

**Figure S4:** Estimated IBD2 and IBD1 probabilities (y-axis) vs. segment length cutoff (x-axis, truffle parameter –S). Model without the inclusion of genotyping error is shown is red, default truffle model is shown in blue. X axis is on log scale.

### 4. Kinship estimation in the 1000 genomes dataset

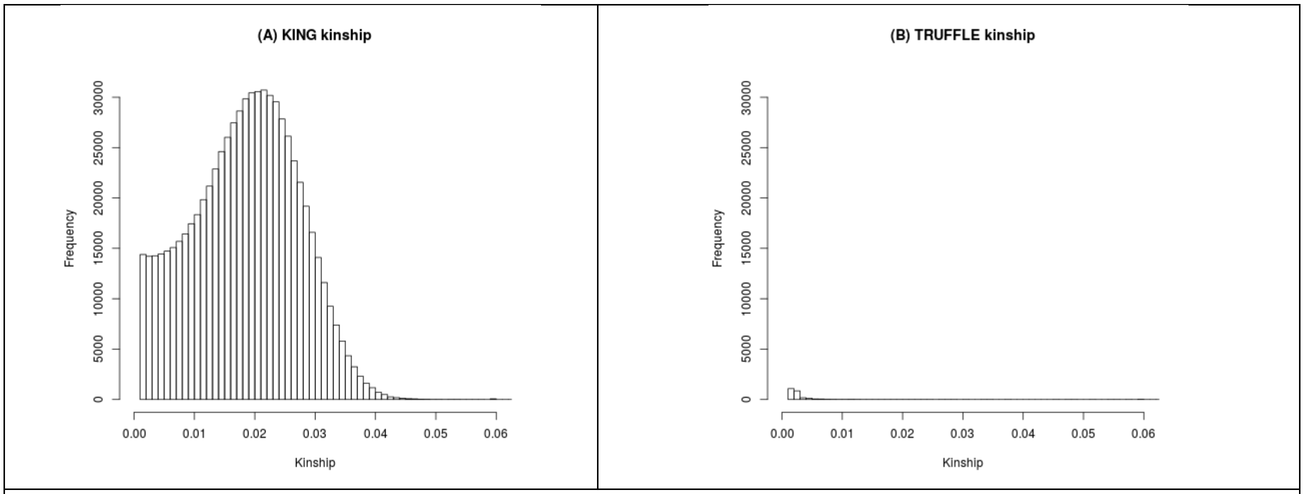


**Figure S5**: Estimation of kinship for 3.1 million pairs in dataset A, shown a histogram of estimated kinships across all pairs. A: KING kinship method shows inflation in kinship estimates below a kinship value of 0.05 (573326 pairs estimated to be second cousin or closer). Dataset A, 65k markers. For clarity pairs with kinship < 0.001 are not shown (2429061 pairs excluded in panel A and 3131206 pairs in panel B). Relationships with kinship >0.06 are also not shown.

**(1) TRUFFLE IBD1 and IBD2**

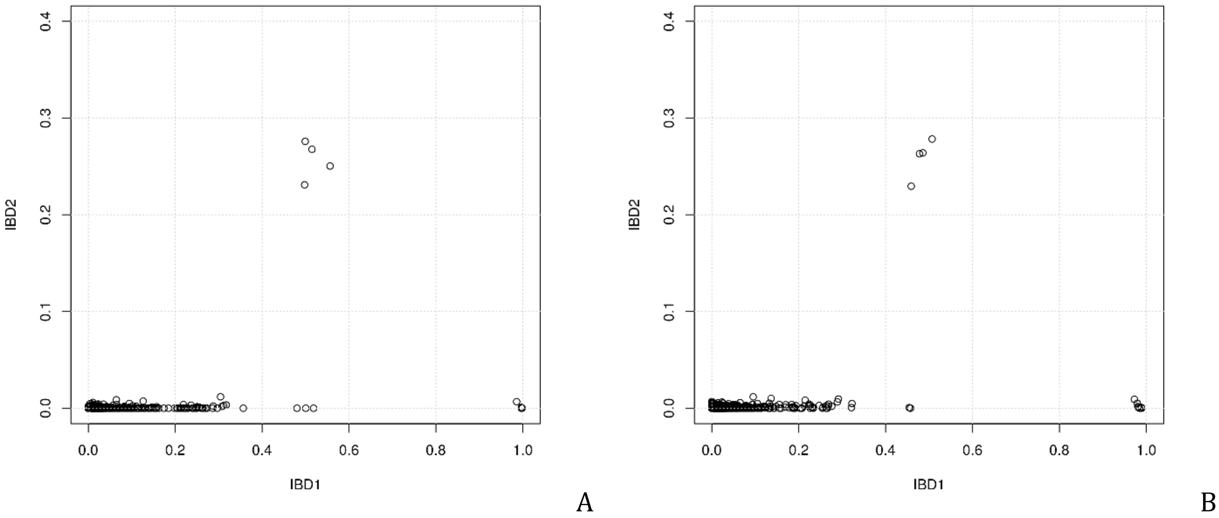


**(2) KING segment IBD1 and IBD2**

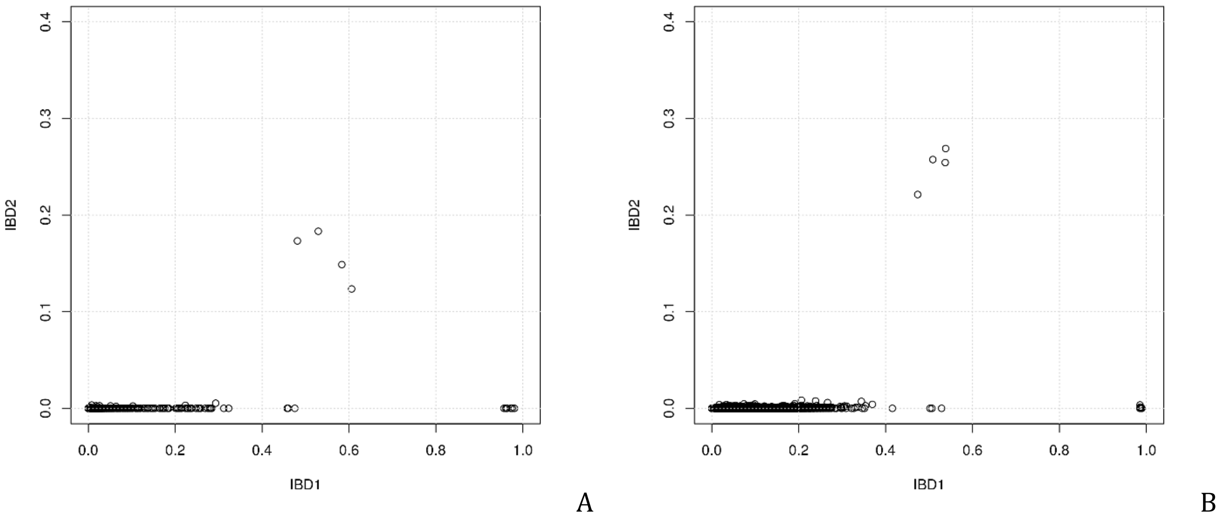


**Figure S6**: IBD1/2 estimation with the truffle and king segment method. (1) TRUFFLE estimation for datasets A and B. (2) KING segment estimation for datasets A and B. The 12 pairs with IBD2 > 0.05 or IBD1 > 0.8 are common in all methods.

### 5. Shared segment analysis by BEAGLE

We downloaded a list of shared segments greater than 5cM among the 2504 individuals in the 1000 genomes dataset, as reported in (Auton, et al., 2015) (Supplementary data section).

These segment lists have been manually curated to join nearby short segments, due to that Refined IBD has no error model and is prone to reporting long segments as multiple short ones.

The data were downloaded from:

[ftp://ftp.1000genomes.ebi.ac.uk/vol1/ftp/release/20130502/supporting/ibd_by_pair/](ftp://ftp.1000genomes.ebi.ac.uk/vol1/ftp/release/20130502/supporting/ibd_by_pair/)

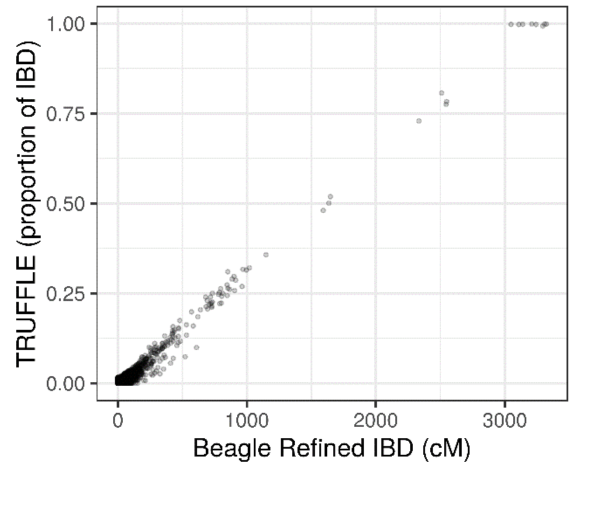


(a) TRUFFLE vs. BEAGLE, correlation = 0.956

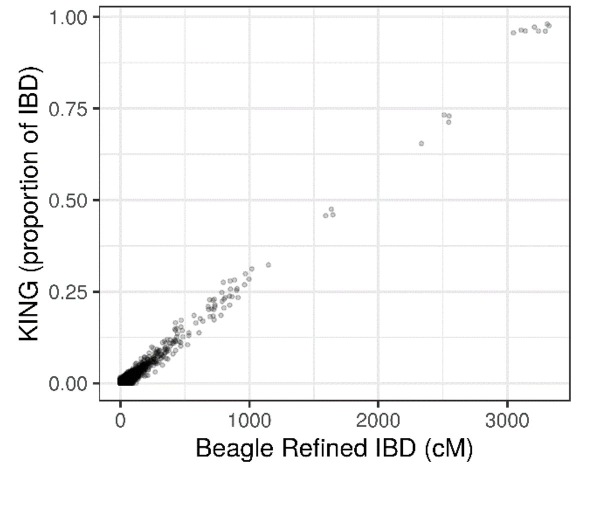


(b) KING vs. BEAGLE correlation = 0.971

**Figure S7:** Comparison of shared segment length identified by (a) TRUFFLE vs. BEAGLE, and (b) KING vs. BEAGLE using the dataset (A) (1000 Genomes data; approx.. 65k markers).

### 6. Kinship estimation comparison between array and sequencing+array data

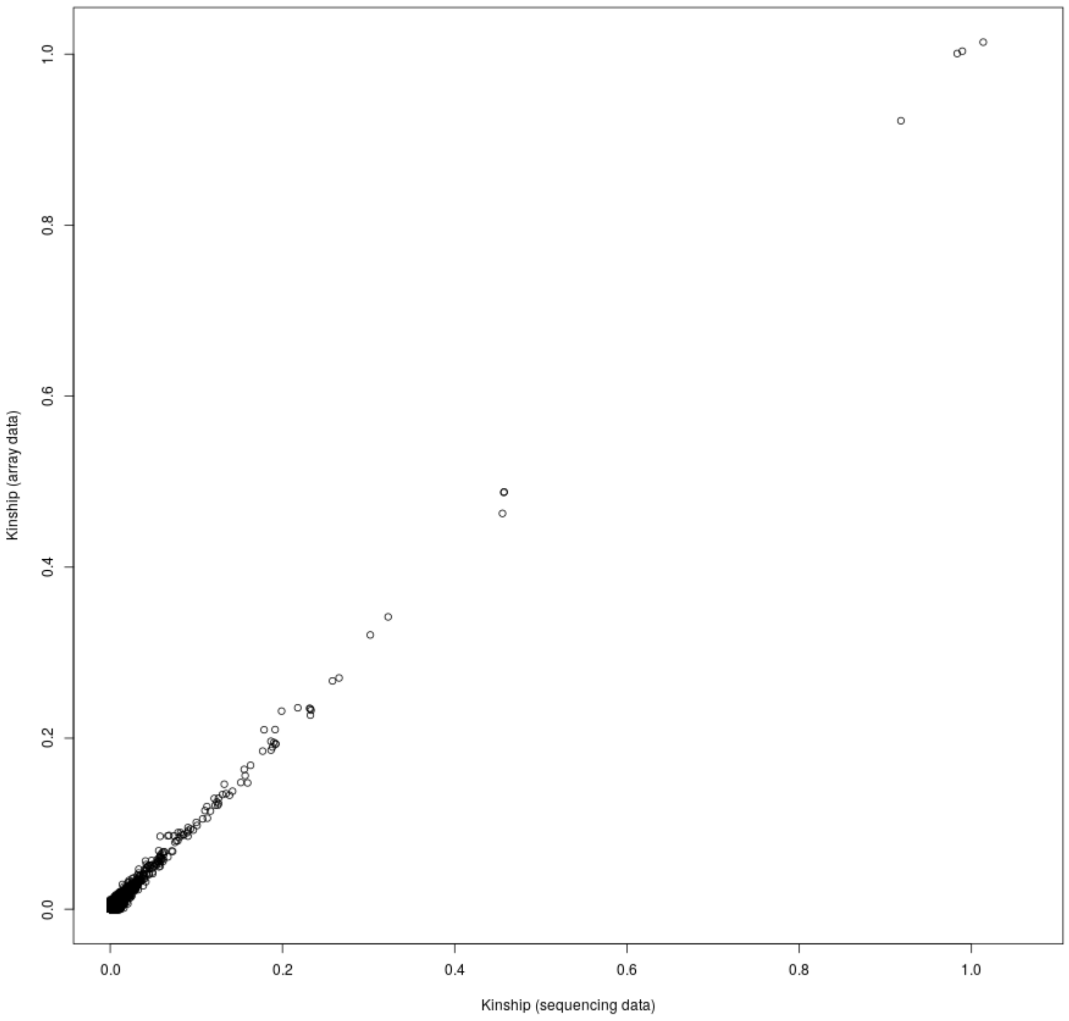


**Figure S8:** Kinship estimation by truffle in 1000 genomes array vs. consensus call data (dataset B). X-axis: sequencing data. Y-axis: array data. Correlation coefficient r = 0.93.

### 7. Identity by Descent analysis within populations

| Population | % of pairs sharing an IBD1 segment of length >5cM | % of pairs sharing an IBD1 segment of length >10cM | % of pairs sharing an IBD2 segment of length >5cM | Number of pairs |
|---|---|---|---|---|
| ACB | 9.17 | 5.60 | 0.000 | 4517 |
| ASW | 2.21 | 0.89 | 0.000 | 1807 |
| BEB | 5.86 | 0.55 | 0.000 | 3649 |
| CDX | 43.26 | 7.54 | 0.000 | 4191 |
| CEU | 12.35 | 0.70 | 0.000 | 4834 |
| CHB | 14.87 | 0.21 | 0.000 | 5253 |
| CHS | 19.27 | 1.66 | 0.000 | 5429 |
| CLM | 61.52 | 40.47 | 0.077 | 3919 |
| ESN | 26.58 | 14.29 | 0.000 | 4710 |
| FIN | 74.72 | 18.48 | 0.021 | 4790 |
| GBR | 19.25 | 5.75 | 0.025 | 4053 |
| GIH | 35.73 | 8.88 | 0.000 | 5181 |
| GWD | 9.01 | 3.19 | 0.016 | 6270 |
| IBS | 10.15 | 0.65 | 0.000 | 5666 |
| ITU | 11.21 | 1.48 | 0.000 | 5138 |
| JPT | 30.36 | 0.67 | 0.000 | 5356 |
| KHV | 24.71 | 1.88 | 0.000 | 4848 |
| LWK | 40.97 | 13.57 | 0.000 | 4681 |
| MSL | 24.07 | 10.97 | 0.000 | 3519 |
| MXL | 38.71 | 5.03 | 0.000 | 2007 |
| PEL | 78.64 | 10.02 | 0.000 | 3562 |
| PJL | 14.61 | 5.18 | 0.022 | 4496 |
| PUR | 82.67 | 62.02 | 0.289 | 4495 |
| STU | 23.41 | 7.90 | 0.000 | 5079 |
| TSI | 21.82 | 7.58 | 0.000 | 5623 |
| YRI | 4.11 | 1.51 | 0.000 | 5773 |

**Figure S9:** For each same population pair we compute the percent of pairs sharing: (1) at least an IBD1 segment of length 5cM, (2) at least an IBD1 segment of length 10cM, (3) at least an IBD2 segment of length 5cM.

### 8. Short segment density plots occurring within populations.

For all the populations in the 1000 genomes dataset, we generate segment density plots that highlight the regions of extended IBD1 segment sharing for every chromosome. The blue rectangles denote gaps in the genome assembly, including centromere regions and other large gaps of more than 1MBp.

**Figure S10:** Distribution and locations of segments shared within population in the 1000 genomes data. Estimated from variant set (B).

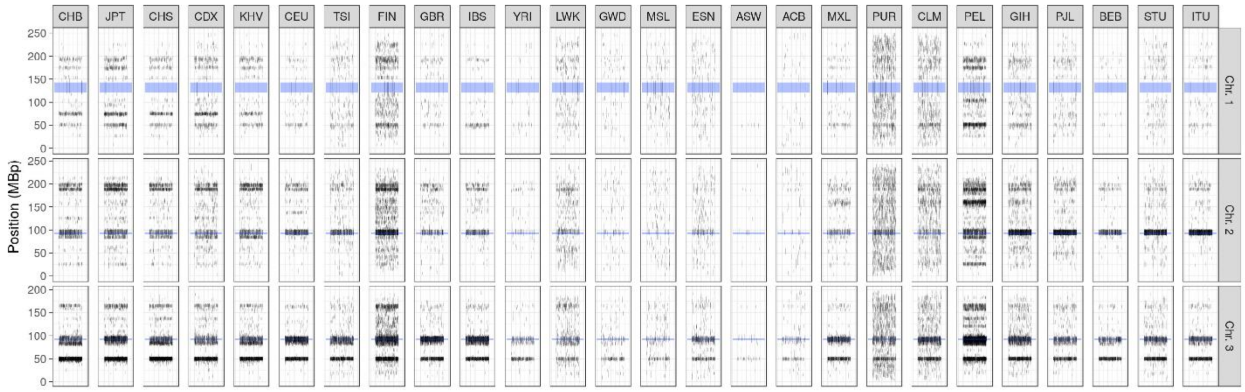


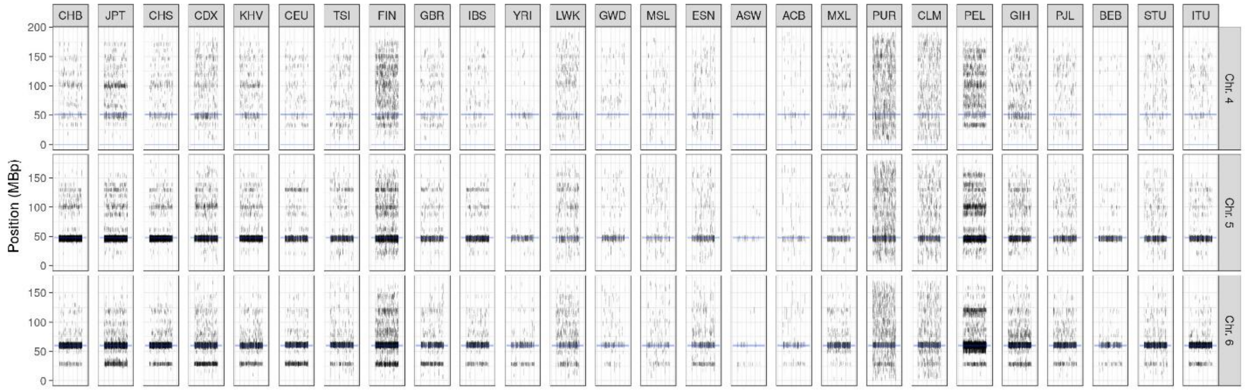

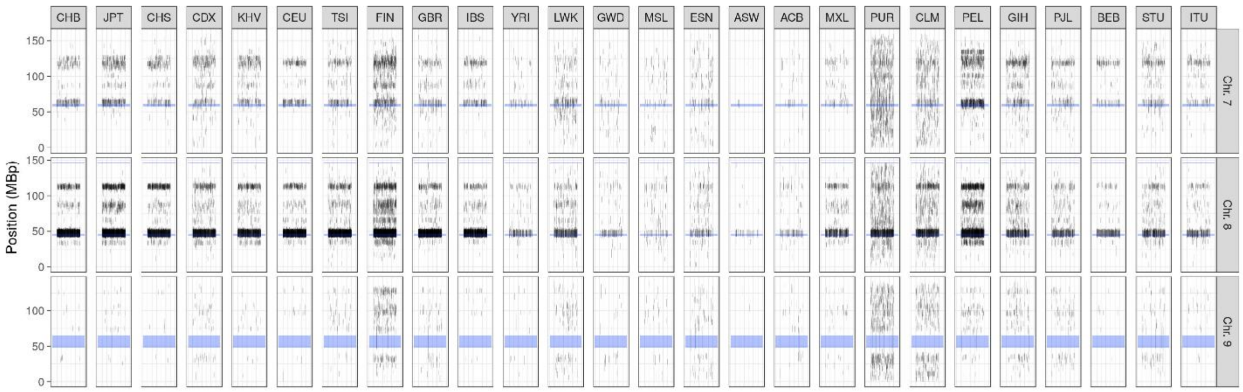
CHB JPT CHS CDX KHV CEU TSI FIN GBR IBS YRI LWK GWD MSL ESN ASW ACB MXL PUR CLM PEL GIH PJL BEB STU ITU
Position (MBp)
Chr. 7
Chr. 8
Chr. 9

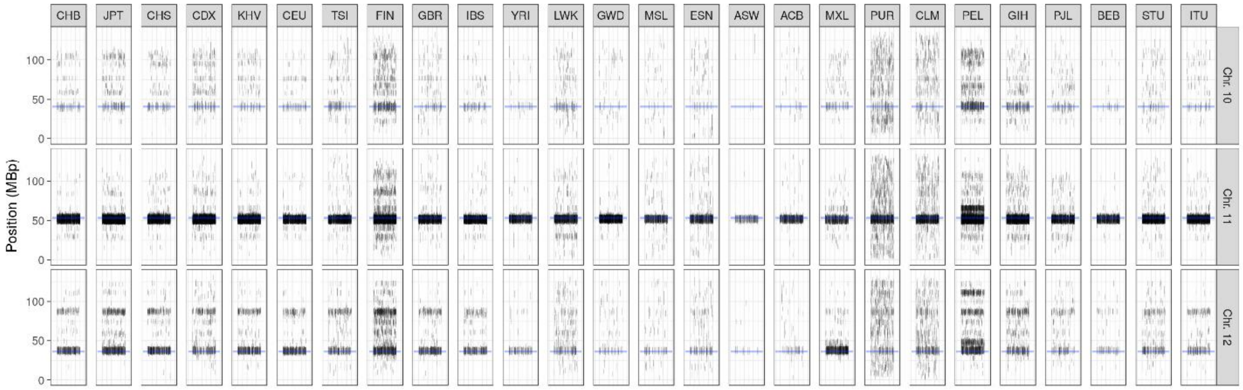
CHB JPT CHS CDX KHV CEU TSI FIN GBR IBS YRI LWK GWD MSL ESN ASW ACB MXL PUR CLM PEL GIH PJL BEB STU ITU
Position (MBp)
Chr. 10
Chr. 11
Chr. 12

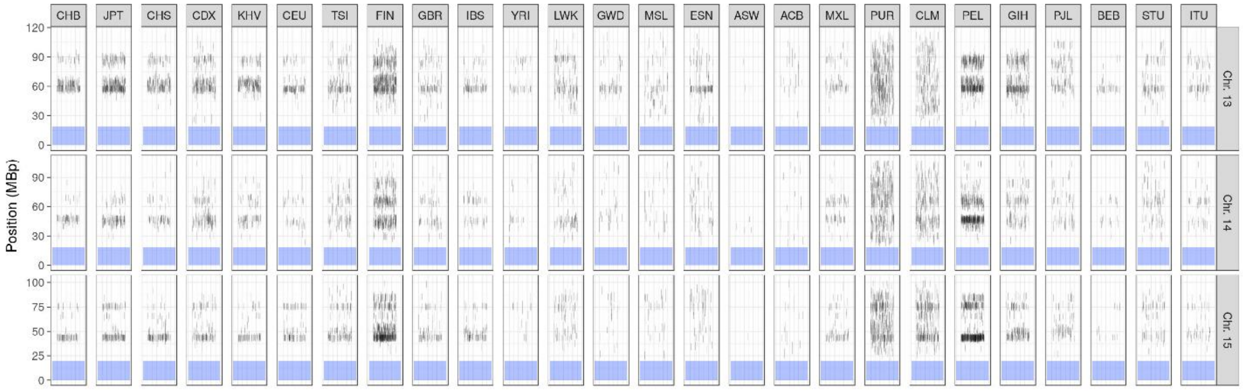
CHB JPT CHS CDX KHV CEU TSI FIN GBR IBS YRI LWK GWD MSL ESN ASW ACB MXL PUR CLM PEL GIH PJL BEB STU ITU
Position (MBp)
Chr. 13
Chr. 14
Chr. 15

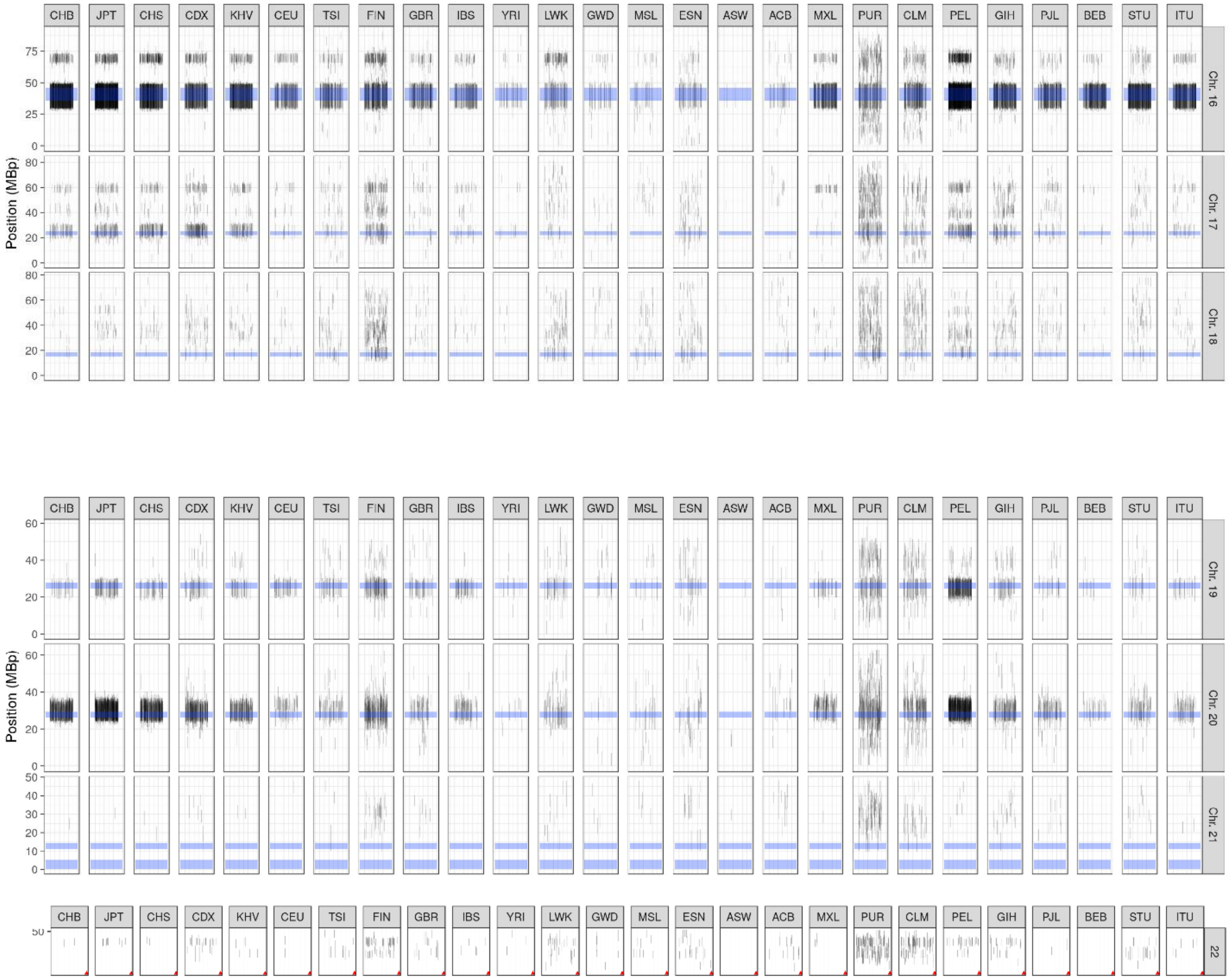

CHB
JPT
CHS
CDX
KHV
CEU
TSI
FIN
GBR
IBS
YRI
LWK
GWD
MSL
ESN
ASW
ACB
MXL
PUR
CLM
PEL
GIH
PJL
BEB
STU
ITU
Position (MBp)
Chr. 16
Chr. 17
Chr. 18
Chr. 19
Chr. 20
Chr. 21
22

### 9. IBD segments in 1000 genomes between different methods and SNPs

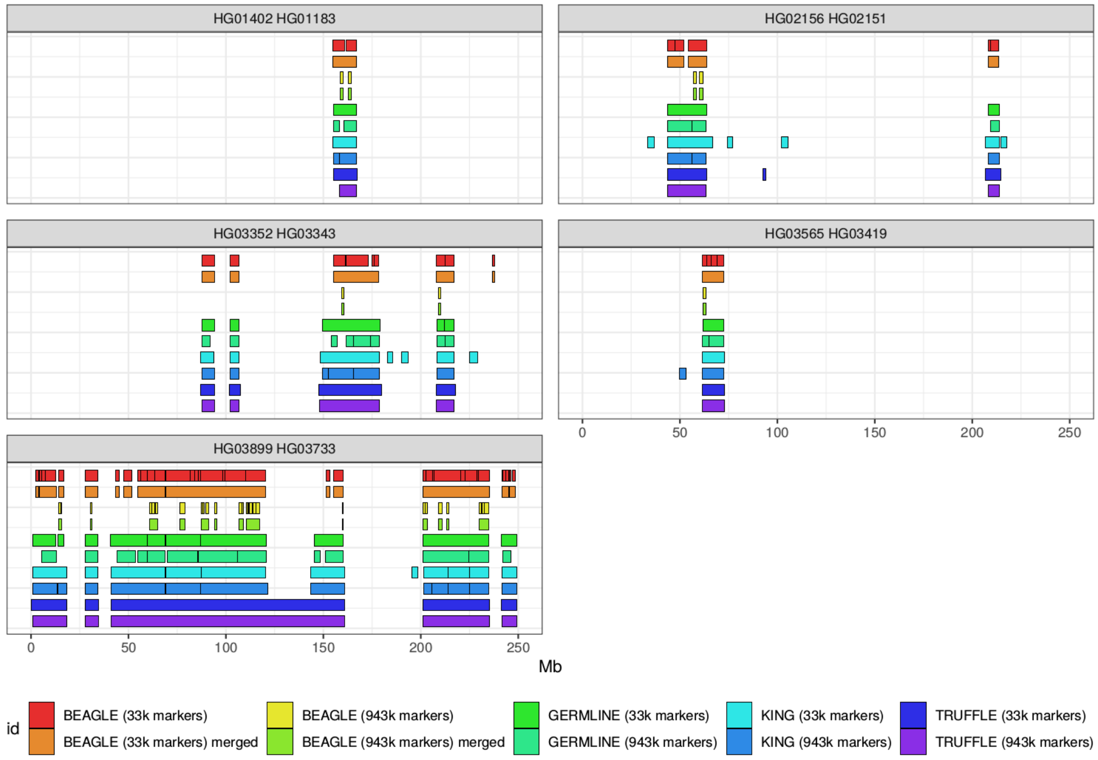


**Supplementary Figure S11. Comparison of locations of IBD segments on chromosome 1 from the 1000 Genomes Project for 5 randomly selected pairs, 1 each from full-sibs and 1st cousins, and 3 more distantly related pairs.** The data are from phase 3 release 5. KING and TRUFFLE can work on unphased data, and BEAGLE Refined IBD and GERMLINE were applied to the data previously phased by the 1000 genomes analysis group using both BEAGLE and Shapeit2. The 33k SNPs have MAF >5% with > 5 kb between two consecutive SNPs with missing rate <2%, and the 943k SNPs have MAF >1% and missing rate <2%. Positions are based on build 37, where the centromere is located at 121.5 - 142.5 Mb.

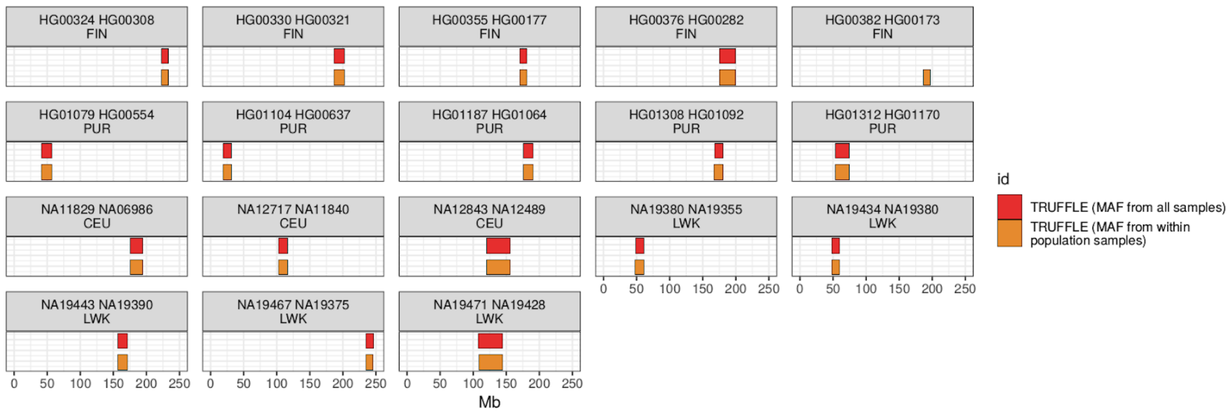


**Supplementary Figure S12. Comparison between segment locations inferred when using global MAF cutoff criteria vs population-specific MAF from Dataset B using TRUFFLE.** Specifically we run TRUFFLE on the CEU, LWK, FIN, and PUR populations of the 1000 Genomes Project, estimating allele frequencies from either the whole 26 populations or within each population analyzed, using (i) the primary dataset B using cross-population MAF >5% and spacing at least 5 kb, and (ii) the comparison dataset B using within-population MAF >5% and spacing at least 5 kb. Presented are 18 randomly selected within-population pairs on chromosome 1 from CEU, LWK, FIN and PUR 1000 Genomes Project populations.

### 10. Kinship estimation in the GAW20 data

| | | Inferred Degree of Relationship | | | | | | |
|---|---|---|---|---|---|---|---|---|
| | | 1 | 2 | 3 | 4 | 5 | 6 | >= 7 |
| Actual Degree of Relationship | 1 | 882 | 0 | 0 | 0 | 0 | 0 | 0 |
| | 2 | 1 | 678 | 10 | 0 | 0 | 0 | 0 |
| | 3 | 0 | 8 | 464 | 50 | 1 | 0 | 0 |
| | 4 | 0 | 0 | 5 | 144 | 48 | 4 | 1 |
| | 5 | 0 | 0 | 0 | 1 | 7 | 9 | 3 |

**Table S1**: Actual vs. estimated degree of relationship in the GAW20 dataset. Pairs with pedigree relationship specified as non-zero as shown. The estimated degree of relationship is computed from the kinship $k$ as the closest integer to $-\log_2 k - 1$.

**Table S2:** Summary of GOLDN analyses by TRUFFLE and KING.

| Degree | Pedigree-based Kinship Coeff. | TRUFFLE estimates | TRUFFLE Std.Dev. | KING-segment 2.1.6 estimates | KING Std.Dev. |
|---|---|---|---|---|---|
| 1 | 0.25 | 0.252 | 0.0174 | 0.251 | 0.0171 |
| 2 | 0.125 | 0.125 | 0.0165 | 0.129 | 0.0155 |
| 3 | 0.0625 | 0.0606 | 0.0144 | 0.069 | 0.0136 |
| 4 | 0.03125 | 0.0272 | 0.00880 | 0.0369 | 0.00872 |
| 5 | 0.015625 | 0.0106 | 0.00470 | 0.021 | 0.00492 |
| >5 | 0 | 0.000107 | 0.00037 | 0.00023 | 0.00155 |

### 11. Simulation scripts used for the power study

The following script was used as a basis for generating an initial population used in the power study. It was adapted from example4 in the simuGWAS collection of population simulation scripts (https://github.com/BoPeng/simuPOP-examples/tree/master/published/simuGWAS).

```
#!/usr/bin/env python

#
import sys, os, logging

from simuOpt import Params, setOptions
#setOptions(alleleType='binary', optimized=False, gui=False)

from simuPOP import *

import loadHapMap3, selectMarkers, simuGWAS


def downloadData(logger):
    '''
    Download and create populations from the third phase of the HapMap3 data.
    This equivalent to command

    > loadHapMap3.py --chroms=2
    '''
    if not os.path.isdir('HapMap'):
        os.mkdir('HapMap')
    for chrom in chroms:
        for popName in loadHapMap3.HapMap3_pops:
            filename = 'HapMap/HapMap3_%s_chr%d.pop' % (popName, chrom)
            if not os.path.isfile(filename):
                pop = loadHapMap3.loadHapMapPop(chrom, popName, logger)
                pop.save(filename)

def getInitPop(logger):
    '''
    Step 2: Select 2000 markers on a random regions on chromosomes 2, with minor allele frequency 0.05.

     > selectMarkers.py --chroms=2 --numMarkers=2000 --startPos=50000000 --filename=ex4_init.pop --
minAF=0.05
       --minDist=50000 --HapMap_pops="['HapMap3_JPT+CHB','HapMap3_CEU']" --mergeSubPops=False
    '''
    if os.path.isfile('ex4_init.pop') and os.path.isfile('ex4_init.pop.lst'):
        if logger:
            logger.info('ex4_init.pop already exists. Please remove this file if you would like to
regenerate an initial population.')
        return
    if logger:
        logger.info('Select 5000 markers from chromosomes 2')
    pop = selectMarkers.getHapMapMarkers(
        HapMap_dir='HapMap',
        chroms=[1],
        HapMap_pops=['HapMap3_TSI','HapMap3_LWK'],


        startPos=[10000000],
        minAF=0.05,
        minDist=2000,
        numMarkers=[25000],
        mergeSubPops=False,
        logger=logger)
    if logger:
        logger.info('Saving initial population to ex4_init.pop')
    pop.save('ex4_init.pop')
```

```
    createFiles(pop, "init")

    if logger:
        logger.info('Saving marker information to ex4_init.pop.lst')
    selectMarkers.saveMarkerList(pop, 'ex4_init.pop.lst', logger)


def expandPop(logger):
    # This is equivalent to
    # > simuGWAS.py --initPop=ex3_init.pop --migrRate=0.0001 --scale=5
    #
    # This just to make this result reproducible.
    getRNG().set(seed=1355)
    #
    filename = 'ex4_expanded.pop'
    if os.path.isfile(filename):
        if logger:
            logger.info('%s already exists. Please remove this file if you would like to regenerate an
expanded population.' % filename)
        return
    else:
        if logger:
            logger.info('Simulating an expanded population %s from ex4_init.pop...' % filename)
    pop = loadPopulation('ex4_init.pop')



    pars = Params(simuGWAS.options, initPop=filename, migrRate=0.0001,
                  recIntensity=100e-8,
                  scale=1,
                  expandSize=15000, expandGen=3000)


    pop = simuGWAS.simuGWAS(pars, pop, logger=logger)
    if logger:
        logger.info('Saving expanded population to ' + filename)
    pop.save(filename)


def mix(logger):
    '''Load expanded population and mix using non-random mating'''
    if logger:
        logger.info('Loading population ex4_expanded.pop and mix')
    pop = loadPopulation('ex4_expanded.pop')
    pop.addInfoFields('ancestry')
    # define two virtual subpopulations by ancestry value
    pop.setVirtualSplitter(InfoSplitter(field='ancestry', cutoff = [0.5]))
    # initialize ancestry
    initInfo(pop, [0]*pop.subPopSize(0) + [1]*pop.subPopSize(1), infoFields='ancestry')
    initSex(pop)
    ops=[ MendelianGenoTransmitter(),
          InheritTagger(mode=MEAN, infoFields='ancestry')
        ]
    pop.evolve(
        preOps = Migrator(rate =[
            [0., 0], [0.05, 0]]),
        matingScheme = HeteroMating(
            matingSchemes=[
                RandomMating(ops=ops),
                RandomMating(subPops=[(0,0)], weight=-0.80, ops=ops),
                RandomMating(subPops=[(0,1)], weight=-0.80, ops=ops)
            ],
        ),
        postOps = PyEval(r"'Generation %d\n' % gen"),
        gen=10,
    )
    # remove the second subpop
    if logger:
        logger.info('Removing MKK subpopulation and save admixed population to ex4_mixed.pop')
```

```
    pop.removeSubPops(1)
    pop.save('ex4_mixed.pop')







def createFiles(pop,inputfileroot):
        """Creates inputdata file for PLINK in biallelic format. Assumes inputdata is phased haplotypes,
one line
        per individual."""
        if pop.ploidy() != 2:
                print "PLINK requires biallelic data!"
                return 0

        numInd = pop.popSize()
        locNames = pop.lociNames()
        numLoc = pop.totNumLoci()
        allInd = pop.genotype()

        markerfilename = inputfileroot + ".ped"
        markerOut = open(markerfilename,'w')
        id_counter = 0
        for ind in pop.individuals():
                geno = ind.genotype()
                hap1 = ['1' if x == 0 else '2' for x in geno[:numLoc]]
                hap2 = ['1' if x == 0 else '2' for x in geno[numLoc:]]
                geno_out = " ".join(["%s %s"%(hap1[i],hap2[i]) for i in xrange(numLoc)])
                if ind.affected():
                        outstring = "case%d 1 0 0 1 2 %s\n"%(id_counter,geno_out)
                else:
                        outstring = "control%d 1 0 0 1 1 %s\n"%(id_counter,geno_out)
                id_counter +=1
                markerOut.write(outstring)
        markerOut.close()


        positionfilename = inputfileroot + ".map"
        positionOut = open(positionfilename,'w')
        for loc in xrange(numLoc):
                positionOutString = "%s\t%s\t0\t%s\n"
%(pop.chromName(pop.chromLocusPair(loc)[0]),locNames[loc],pop.locusPos(loc))
                positionOut.write(positionOutString)
        positionOut.close()
        return [markerfilename,positionfilename]






if __name__ == '__main__':
    logging.basicConfig(level=logging.DEBUG)
    logger = logging.getLogger('example4')
   # downloadData([2], logger)
    getInitPop(logger)
    expandPop(logger)

    pop = loadPopulation('ex4_expanded.pop')
    createFiles(pop, "g3000")


    #mix(logger)

    #pop = loadPopulation('ex4_mixed.pop')
    #createFiles(pop, 'ex4_mixed')
```
